# Deletion of the gene for a cyanobacterial ribosome-associated protein affects the carbon/nitrogen metabolism

**DOI:** 10.64898/2026.08.10.743958

**Authors:** Nouha Abdelaziz, Alexander Kraus, Stefan Timm, Friedel Drepper, Viktoria Reimann, Marc Broghammer, Bettina Knapp, Antonio López-Lozano, Ravi Shankar Ojha, Bettina Siebers, Michael Y. Galperin, José Manuel García-Fernández, Manuel Brenes-Alvarez, Pitter F. Huesgen, Martin Hagemann, Wolfgang R. Hess

**Author notes:** **Correspondence:** Wolfgang R. Hess. Co-sharing first authors (Nouha Abdelaziz, Alexander Kraus).

## Abstract

In contrast to their important structural and regulatory functions, such as in the metabolism of cyanobacteria, genes encoding small proteins are often not well characterized. Cyanobacteria use redox equivalents and energy from oxygenic photosynthesis to produce organic carbon compounds from inorganic carbon (C_i_) and organic nitrogen compounds from inorganic nitrogen sources. Therefore, the assimilation and metabolism of carbon and nitrogen are coordinated at multiple levels in cyanobacteria. Here, we analyzed the *Synechocystis* sp. PCC 6803 gene *ssr3189* encoding a 55 amino acids protein. Orthologs were detected in 665 cyanobacterial genomes defining COG5794 in the Database of Clusters of Orthologous Genes. Homologs in several eukaryotic algae suggest that Ssr3189 is an important protein that originated in cyanobacteria, was retained in algae after endosymbiosis, but was lost in plants. Polynucleotide kinase assays validated Ssr3189 as an RNA-binding protein. Deletion of *ssr3189* resulted in lower pigmentation, delayed growth, and alterations in the expression of genes encoding transporters for nitrogen and C_i_, and metabolic enzymes. Metabolomic analysis revealed a substantial overaccumulation of glutamine and tricarboxylic acid cycle intermediates in the deletion mutant, and further differences in the amino acid and organic acid pools compared to the wild type. Co-immunoprecipitation analysis yielded ribosomal protein S21, enolase and the Cas6-1 endoribonuclease as the most strongly co-enriched proteins, together with all other ribosomal proteins and a small set of metabolic enzymes. These findings are consistent with observations that *ssr3189* encodes the ribosome-associated protein cS24 and suggest that it connects translation with metabolic control, and, potentially, RNA decay.

**IMPACT STATEMENT:** Despite considerable progress in analyzing microbial genomes, there are still substantial numbers of uncharacterized gene functions. Here, we analyzed a mutant lacking gene *ssr3189* that is widely conserved, but phenotypically uncharacterized in cyanobacteria. This gene is important for growth at the optimum temperature and essential at lower temperatures. In its absence, important metabolites were overaccumulated, while genes involved in nitrogen and C_i_ uptake were dysregulated. The encoded protein binds RNA and interacts with proteins involved in translation and metabolism. The findings are consistent with a function as a ribosomal protein bridging protein synthesis and the regulation of metabolism.

## INTRODUCTION

Cyanobacteria are the sole prokaryotes that assimilate organic compounds from inorganic carbon (C_i_) through oxygenic photosynthesis (Bryant and Frigaard, 2006), rendering them ecologically highly relevant. They are the main constituents of the vast marine picophytoplankton communities (Partensky et al., 1999; Flombaum et al., 2013), typical photoautotrophs in freshwater ecosystems (Cabello-Yeves et al., 2022) and occur in harsh environment such as thermal springs (Finsinger et al., 2008) and desert biological soil crusts (Xu et al., 2021). Consequently, cyanobacteria are a morphologically and physiologically diverse group of prokaryotes with remarkable stress acclimation capabilities (Stanier and Cohen-Bazire, 1977; Ward et al., 2012; Gaysina et al., 2019). There are several laboratory models for the work with cyanobacteria. A well-established unicellular model is *Synechocystis* sp. PCC 6803 (from here *Synechocystis* 6803), which can be grown under a variety of conditions and easily manipulated (Doello et al., 2026).

There are 83 annotated genes in *Synechocystis* 6803 encoding proteins of 60 amino acids (aa) or less, and 54 of these genes are currently lacking a known function (Kraus and Hess, 2025). Moreover, mounting evidence suggests that many more small protein-coding genes are located in seemingly empty intergenic regions, at unconventional sites within transcripts previously considered non-coding RNAs, or out-of-frame within annotated genes (Hadjeras et al., 2025). Several recently discovered small proteins control critical steps in cyanobacterial metabolism, especially in the coordination of carbon and nitrogen (C/N) metabolism. Examples discovered in *Synechocystis* 6803 include the 52 aa small protein PirA, which interacts with the P_II_ protein and regulates the flux into the ornithine-ammonia cycle (Bolay et al., 2021), the 31 aa peptide SniP3 regulating glyceraldehyde-3-phosphate dehydrogenase (Gap2) activity and possibly further enzymes (Alvarenga-Lucius et al., 2026), or the 44 aa AcnSP interacting with the citrate cycle enzyme aconitase (aconitate hydratase B, AcnB) (de Alvarenga et al., 2020).

Previous GradR analyses in *Nostoc* sp. PCC 7120 identified a 54 aa small protein, Asl3888, as an RNA-binding protein (Brenes-Álvarez et al., 2025). In this work we characterized the homologous gene in *Synechocystis* 6803, *ssr3189*. Using polynucleotide kinase (PNK) assays we demonstrate that Ssr3189 is an RNA-binding protein. The deletion of *ssr3189* led to phenotypical changes, distorted mRNA levels of proteins involved in inorganic carbon (C_i_) and nitrogen uptake, significantly enhanced glutamine synthetase activity, and deviations in the concentrations of key metabolites of these pathways. Co-immunoprecipitation (co-IP) experiments with a C-terminally FLAG-tagged protein revealed the ribosomal protein S21, the enzyme enolase and the CRISPR endoribonuclease Cas6-1 as most strongly co-enriched proteins, together with all other ribosomal proteins, several ribosome-associated factors, the enzymes arginine decarboxylase 1 and 2, delta-aminolevulinic acid dehydratase (HemB), pyruvate dehydrogenase subunit alpha (PdhA), and aconitase (AcnB). The findings point a possible mechanism in which Ssr3189 influences the differential expression and activity of metabolically relevant proteins.

## RESULTS

### A Family of Small Proteins Conserved in Cyanobacteria: Defining COG5794

In *Synechocystis* 6803, the gene *ssr3189* is located on the forward strand, downstream of the *trpS* gene that encodes tryptophanyl-tRNA synthetase and upstream of two genes, *apaH* (*slr1885)* encoding a potential bis(5’-nucleosyl)-tetraphosphatase (symmetrical) and *rsfS* (*slr1886)* encoding the ribosomal silencing factor RsfS (**Figure 1A**). This arrangement is not conserved. Synteny analysis revealed that the most frequent gene upstream is an RNA recognition motif (RRM)-domain RNA-binding protein located in the same orientation as *ssr3189* (**Figure 1B, Figure S1**). The presence of several glycine residues in these proteins following the RRM domain indicated that they belong to the group represented by the glycine-rich, cold-inducible Rbp1 protein in *Synechocystis* 6803 (Tan et al., 2011).

**Figure 1.**
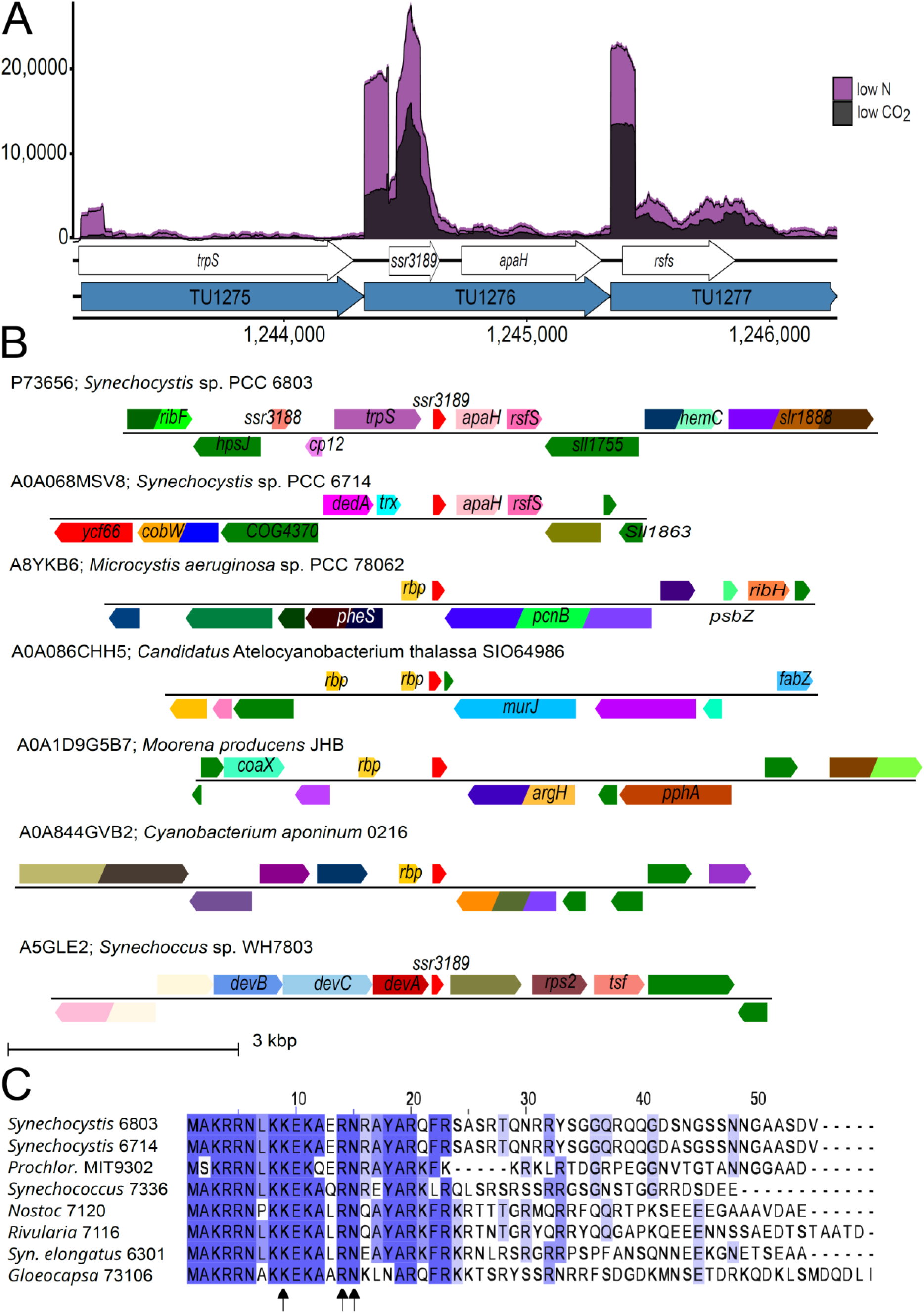
Genomic location and expression of *ssr3189* in *Synechocystis* 6803. **A.** Location of genes *trpS*, *ssr3189*, *apaH* and *rsfs* (white arrows) and the respective transcriptional units TU1275, TU1277 and TU1276 in the chromosome of *Synechocystis* 6803. The transcriptome coverage at low CO_2_ (black) versus nitrogen starvation (purple) is shown. Transcriptional units (TUs) are indicated according to the previous annotation of the transcriptome and genome-wide mapping of transcriptional start sites as blue arrows (Kopf et al., 2014). **B.** Synteny analysis was performed using the EFI Genome Neighborhood Tool (Oberg et al., 2023). Respective gene names are given, likely homologs are in identical colors (red, *ssr3189* and homologs; yellow, homologs of the gene encoding RRM domain protein Rbp1; rose, *apaH*; purple, *trpS*). **C.** Alignment of selected Ssr3189 homologs from cyanobacteria belonging to different morphological sections (Rippka et al., 1979). The alignment is colored according to the percentage identity from low (no shading) to high (blue shading). The residues K9, R14 and N15 conserved in all identified 665 cyanobacterial homologs (**Supplemental Dataset 1**) are marked by arrows. The alignment was generated using ClustalW and visualized by Jalview. An alignment including homologs from cyanobacterial endosymbionts of eukaryotes and from eight major groups of eukaryotic algae, where the homologs are nuclear-encoded and possess an N-terminal targeting sequence, is shown in **Figure S2.**

The *ssr3189* gene encodes a small protein of 55 aa, with a calculated mass of 6.22 kDa and an isoelectric point of 10.7. BlastP database searches identified 665 potential cyanobacterial homologs within strains from all five morphological sections (Rippka et al., 1979). Homologs were also found in the thylakoid-lacking *Gloeobacter violaceus*, the marine *Trichodesmium erythraeum* IMS101, the picocyanobacterial genera *Synechococcus* and *Prochlorococcus*, in thermophilic strains like *Thermosynechococcus sichuanensis*, and in the very streamlined genomes of *Atelocyanobacterium thalassa* or symbionts like *Calothrix rhizosoleniae* and the *Braarudosphaera bigelowii* nitroplast (**Supplementary Dataset 1**). The lengths of these proteins vary from 46 aa in *Gloeomargarita lithophora* (Genbank entry WP_172819669) to 88 residues in *Trichormus azollae* (Genbank entry WP_347611212).

An investigation of the Database of Clusters of Orthologous Genes (COG (Galperin et al., 2025)) revealed that homologs of the *ssr3189* gene were present in 67 out of 75 reference genomes, leading to the definition of COG5794 (**Supplementary Dataset 2**). In two strains, *Stanieria cyanosphaera* PCC 7437 and *Gloeothece verrucosa* PCC 7822, two gene copies were identified. These gene copies are non-identical. For example, the proteins encoded by genes Sta7437_3204 and Sta7437_1591 in *S. cyanosphaera* PCC 7437 show a sequence identity of 56%.

Homologs of COG5794 were not identified in a small number of cyanobacteria, including *Planktothrix agardhii* No.976 and the ten other genomes available for the genus *Planktothrix*, in *Picosynechococcus* sp. PCC 7002 (and 7003), and in bacteria outside of the phylum cyanobacteria (**Supplementary Dataset 2**). However, homologous genes exist in the more recently acquired plastid genomes of the Cercozoa *Paulinella longichromatophora* (AUG32087.1), *Paulinella micropora* (YP_009530514) and *Paulinella chromatophora* (YP_002048915.1), and in the nuclear genomes of Pseudoscourfieldiophyceae green alga (*Pycnococcus provasolii*, XRA98681.1; *Pseudoscourfieldia marina*, XRB13187.1), the haptophyte *Emiliania huxleyi* CCMP1516 (XP_005771629.1) and several red and brown algae (**Figure S2**). Examination of protein sequence alignments revealed striking conservation in the region matching the first 23 amino acids, with K9, R14, and N15 exhibiting complete conservation across all 665 homologs (**Figure 1C**). The protein’s very strong alkaline pI is caused by the fact that about 25% of the protein, especially the conserved N-terminal region, consists of basic amino acids, of which arginine (18%) and lysine (7%) are the most abundant amino acids.

The *ssr3189* gene is part of the transcriptional unit (TU) TU1276, starting with a transcriptional start site at position 1244330, followed by a 108 nt 5’UTR (Kopf et al., 2014). The transcription of TU1276 previously was found to be induced by shifts to nitrogen deprivation and repressed in the absence of C_i_ (**Figure 1A**).

In order to investigate the possible functional significance of Ssr3189, a series of strains were created (**Figure S3A**): A gene deletion strain Δ*ssr3189*; the complementation strain Δ*ssr3189*::*ssr3189*×3F, in which transcription occurs from the native promoter, and an overexpression strain under control of the copper-inducible P*_petE_* promoter on plasmid pVZ322 (strain P*_petE_*-*ssr3189*×3F). In both the complementation and overexpression strains, the coding sequence of *ssr3189* was fused to a sequence encoding a C-terminal 3×FLAG tag. The respective genotypes were verified by PCR (**Figure S3B,C**). Transcription of *ssr3189* in the wild type and Δ*ssr3189*::*ssr3189*×3F was tested by Northern hybridization (**Figure S3D**), showing in both lines a slight increase in mRNA levels under nitrogen deprivation as previously observed (Kopf et al., 2014), confirming that the native promoter had been cloned correctly.

In Western blot analysis using a FLAG tag-specific antibody, a strong ∼15 kDa signal was observed in Δ*ssr3189*::*ssr3189*×3F in extracts from cells collected from stationary phase, standard growth conditions, and nitrogen starvation for 3, 6 and 24 h (**Figure 2A**). With a calculated molecular mass of 8.95 kDa (6.22 kDa for Ssr3189 and 2.73 kDa for the FLAG epitope), the 15 kDa signal could be related to an artefact of resolving small proteins in conventional SDS gels. However, the presence of additional bands corresponding to higher molecular masses suggested potential stable interactions with other proteins. Among these, the signal at 35 kDa was the strongest. These findings indicated that the gene product of *ssr3189* is a rather abundant small protein that may exhibit strong interactions with unknown interacting proteins.

**Figure 2.**
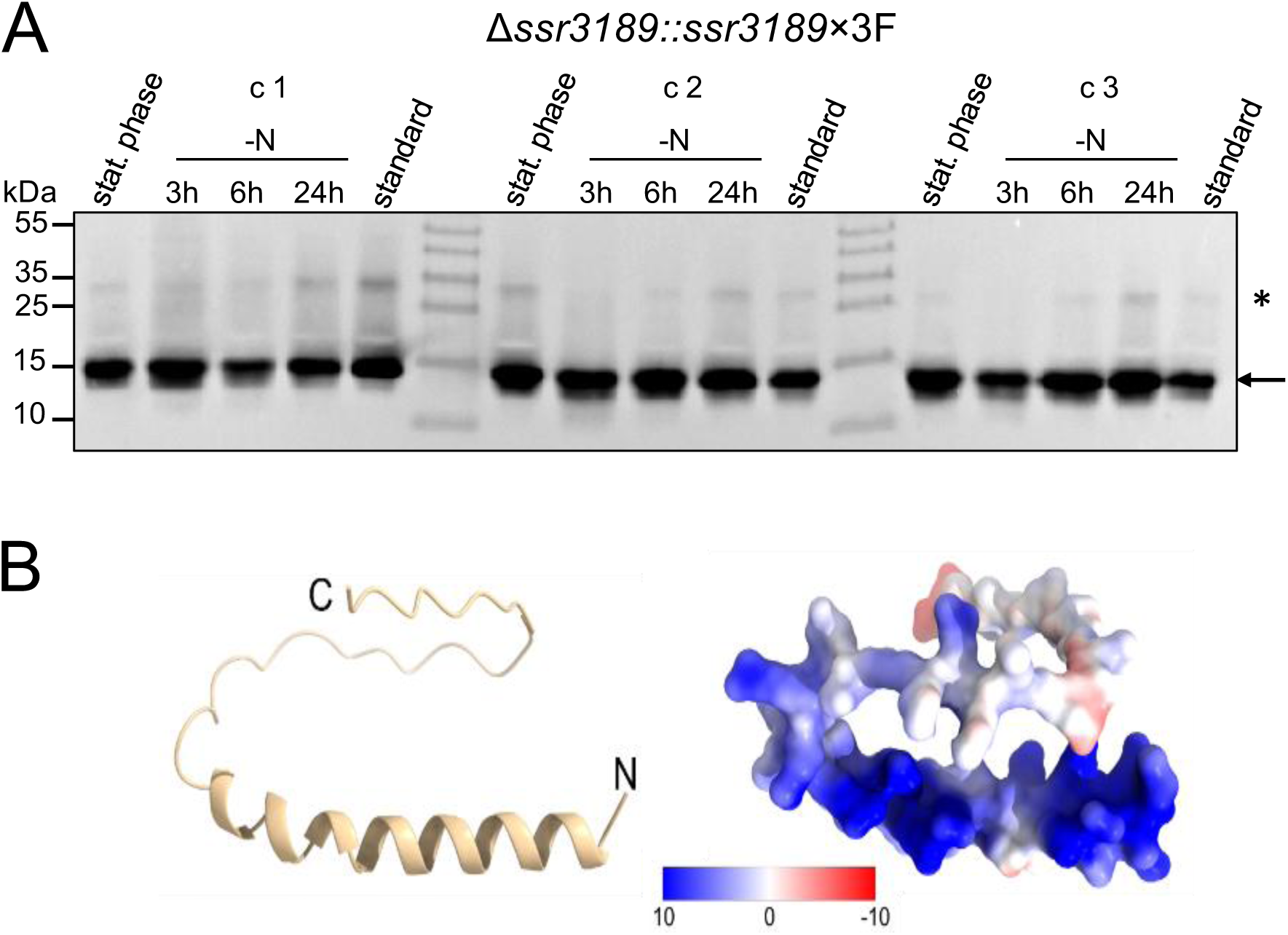
Expression and structure of Ssr3189. **A.** Ssr3189 accumulates as an abundant protein with an apparent molecular mass of 15 kDa (arrow) under three different conditions (stationary phase, standard growth conditions, and nitrogen starvation for 3, 6 and 24 h). A weaker additional signal at 35 kDa (asterisk) was reproducibly observed. Tagged Ssr3189 under the control of its native promoter was expressed in the background of the deletion mutant (strain Δ*ssr3189*::*ssr3189*x3F) and detected by Western blotting using anti-FLAG antiserum at a titer of 1: 5,000. Three biological replicates (c1, c2 and c3) were analyzed. Prestained PageRuler^TM^ (Thermo Fisher Scientific) was used as a molecular mass marker. **B.** *Synechocystis* 6803 Ssr3189 protein structure predicted by AlphaFold (Jumper et al., 2021; Mirdita et al., 2022), indicating the presence of an α-helix in the most conserved part and an intrinsic disordered C-terminal tail. Electrostatic surface representation of Ssr3189 highlighting the electropositive (blue) surface of the α-helix. C- and N-termini are indicated for orientation.

The AlphaFold prediction identified a single α-helix within the first 26 amino acids — but no evidence for a possible membrane association — and a disordered region in the rest of the sequence (**Figure 2B**). We conclude that *ssr3189* and its homologs are widely conserved throughout the cyanobacterial phylum and encode a small basic protein.

### Deletion of *ssr3189* Leads to Phenotypical Changes

Given its regulated expression, the fact that homologs exist in most, but not all cyanobacteria, and not outside the cyanobacterial phylum, we wondered about the possible functional consequences of an *ssr3189* deletion. Indeed, the deletion of *ssr3189* was under laboratory standard growth conditions not lethal, but led to changes in pigmentation and growth (**Figure 3**). In liquid cultures, doubling times reached 7.7 h for the wild type, but 12.5 h for Δ*ssr3189* (**Supplementary Dataset 3**). Interestingly, on plates incubated at 20 °C, no colonies developed for the Δ*ssr3189* mutant, while numerous colonies appeared for the wild type up to a dilution of 10^-3^ (**Figure 3C**). However, the growth effect of the *ssr3189* deletion was much weaker at the optimum growth temperature of 30 °C (**Figure S4**). The complementation strain Δ*ssr3189*::*ssr3189*×3F did grow at 30 °C similar to wild type in liquid cultures (**Figure 3A**) and on plates (**Figure S4**). At 20 °C, strain Δ*ssr3189*::*ssr3189*×3F did grow, although not as well as the wild type, while the overexpression strain P*_petE_*::*ssr3189*×3F grew similar to the wild type (**Figure 3C**).

**Figure 3.**
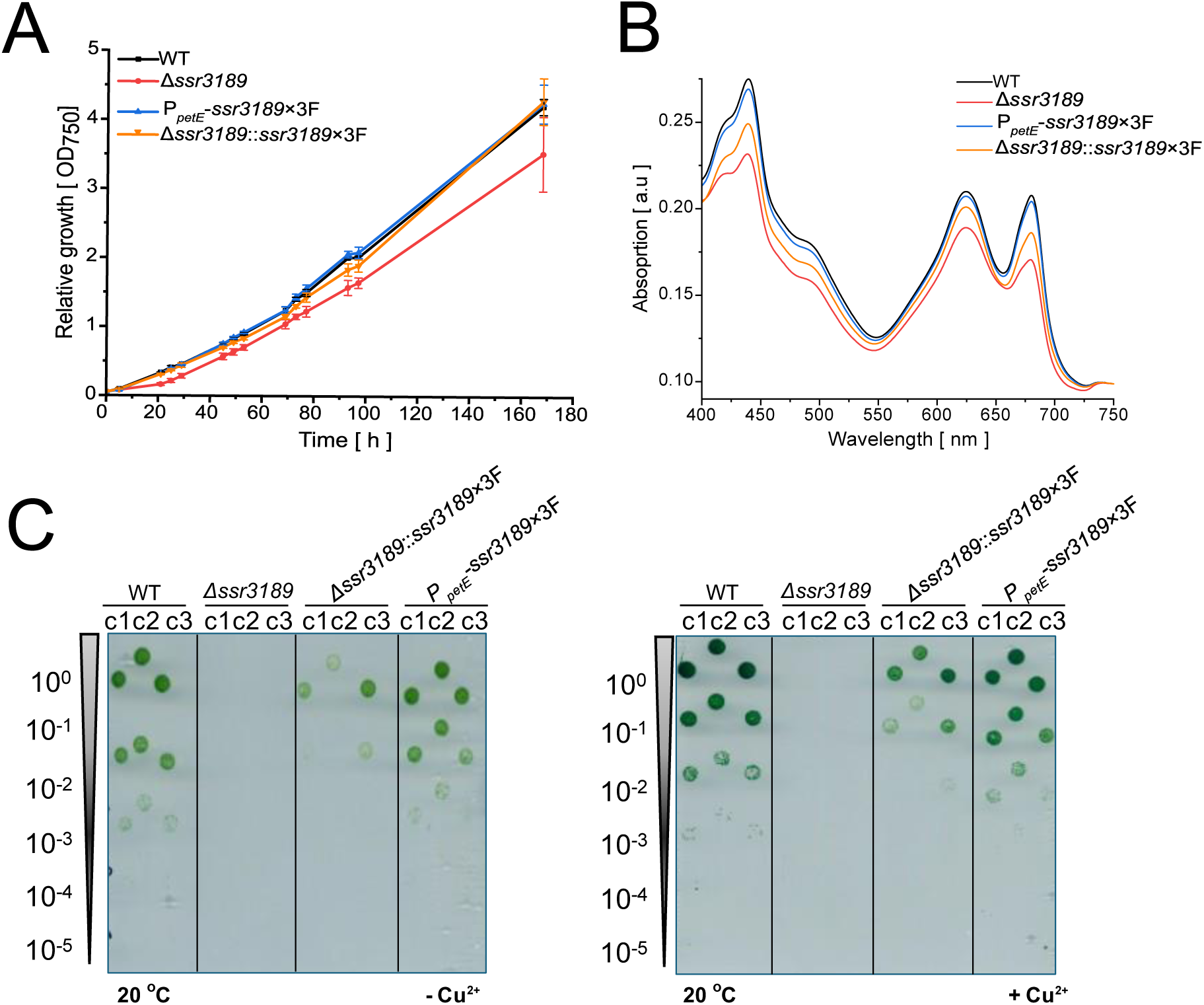
Phenotypical differences between *ssr3189* mutant strains and the wild type. **A.** Growth of wild type (black), the Δ*ssr3189* deletion mutant (red), the *ssr3189* overexpression strain under the control of the P*_petE_* promoter in the wild type background (blue, P*_petE_*-*ssr3189*×3F), and the complementation strain expressing *ssr3189* under the control of the P*_ssr3189_* promoter in the Δ*ssr3189* background (orange, Δ*ssr3189*::*ssr3189*×3F). Data points represent the average of 3 biological replicates. Details in **Supplemental Dataset 3**. **B.** Room temperature absorption spectra for wild-type and the indicated *ssr3189* mutant strains normalized to wild-type OD_750_ (same strains and colors as in panel (A)). Spectra were recorded from three biological replicates and averaged. **C.** Left panel: Drop dilution assay on solid medium without copper in biological triplicates, c1 to c3, of the *Synechocystis* 6803 wild type (WT), the *ssr3189* deletion mutant Δ*ssr3189,* the complementation and overexpression strains Δ*ssr3189*::*ssr3189*×3F and P*_petE_*-*ssr3189*×3F, under continuous light of 40 µmol photons m^-2^ s^-1^, at 20 °C. Strains were pre-cultivated in liquid BG11 medium, P*_petE_*-*ssr3189*×3F was pre-cultivated in BG11 without CuSO_4_, under constant light at 30 °C. The indicated different dilutions were spotted. Right panel: BG11 medium containing 0.3 µM CuSO_4_ was used. Plates were photographed after 15 d of incubation.

Room temperature absorption spectra showed that the phycobilisome peak at approximately 625 nm, as well as the chlorophyll peaks at 440 and 680 nm were decreased in the Δ*ssr3189* mutant compared to wild type. The complementation strain Δ*ssr3189::ssr3189*×3F did not fully recover the phenotype in pigmentation, although the peak intensities in the absorption spectra were clearly higher than in Δ*ssr3189* (**Figure 3B**). The overexpression strain P*_petE_*-*ssr3189*×3F showed a spectrum indistinguishable from wild type (**Figure 3B**).

In summary, the data demonstrated that the protein encoded by *ssr3189* was essential for cell viability at 20 °C, but not at 30 °C, although it was still required for optimal growth. The complete or partial complementation observed for the complementation and overexpressor strains indicated that the protein maintained its functionality regardless of the C-terminally fused 3×FLAG epitope. The observed phenotypic effects in Δ*ssr3189* suggested a critical role in cell vitality.

### Transcriptional Dysregulation of Nitrogen and Inorganic Carbon Transporter Genes in the Δ*ssr3189* Mutant

Transcriptomic analyses were performed to study if the phenotypic effects associated with the absence of *ssr3189* were linked to changed transcript accumulation. We used microarrays that cover all transcripts identified in thorough transcriptome analyses (Kopf et al., 2014) and which enable the direct detection of transcripts from total RNA preparations, without reverse transcription (Voß and Hess, 2014). Transcript levels were determined in wild type and Δ*ssr3189* strains cultivated under standard conditions (**Supplemental Dataset 4**). Eighteen genes were up-and 64 genes were downregulated in the absence of *ssr3189* compared to wild type at standard conditions (|log_2_FC| ≥ 0.6, p value ≤ 0.05) (**Figure 4A**). The up-regulated genes in Δ*ssr3189* were mainly associated with C_i_ uptake, especially *cmpA* and *cmpB* (Omata et al., 1999), including the operon regulator CmpR (Nishimura et al., 2008) (**Figure 4A, B**). Further up-regulated genes encode the Flv2-Sll0218 photoprotective system (Bersanini et al., 2017), and the alternative linker protein ApcI involved in remodeling phycobilisome structures under different light conditions and upon chemically induced reduction of the plastoquinone pool (Espinoza-Corral et al., 2025).

**Figure 4.**
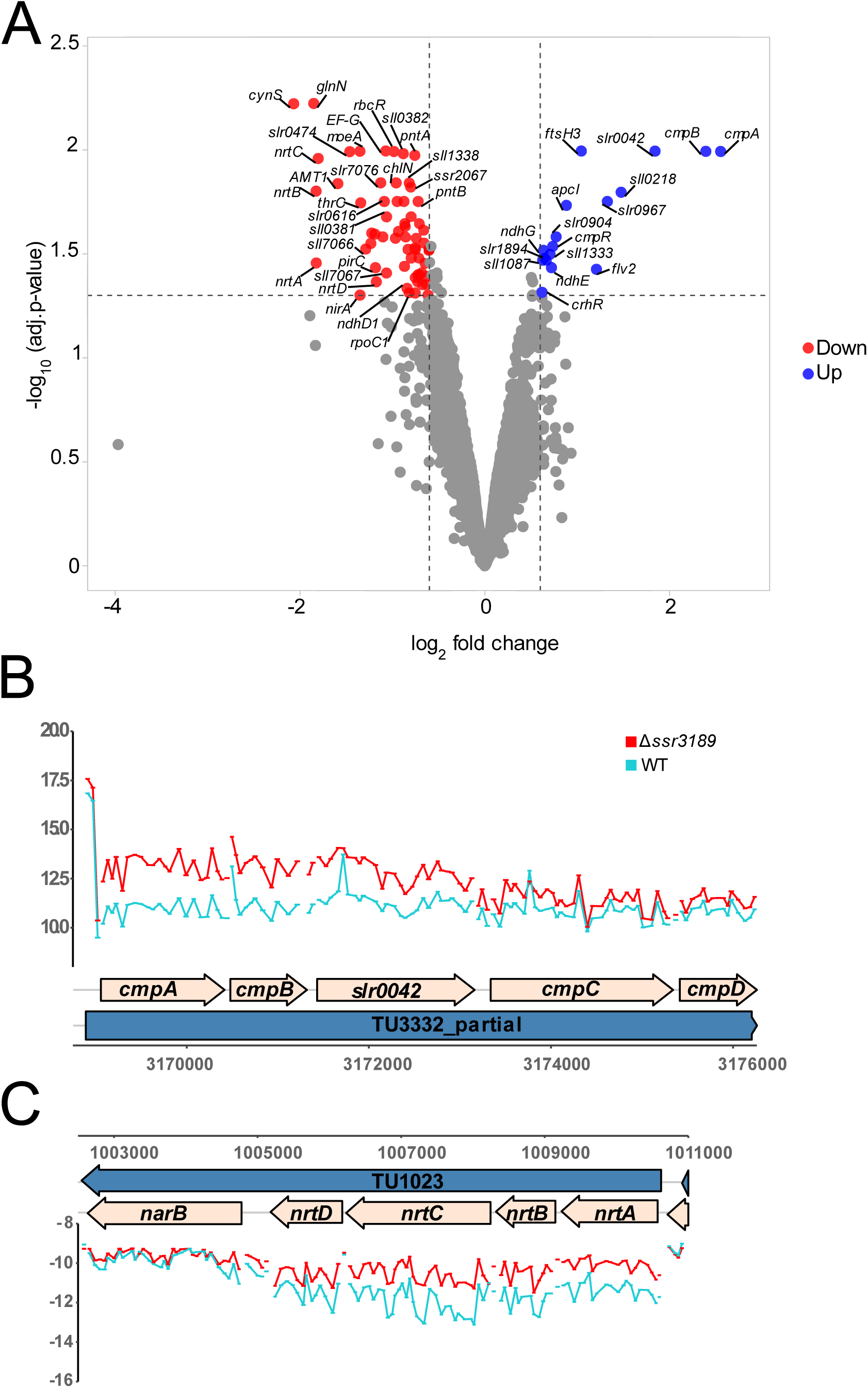
Deletion of *ssr3189* leads to transcriptional changes in genes encoding proteins of the nitrogen and carbon transporter complexes in *Synechocystis* 6803. **A.** Volcano plot of microarray analysis of wild type compared to Δ*ssr3189* under standard conditions. Upregulated genes are shown in blue and downregulated genes in red. Thresholds were set at |log_2_FC| ≥ 0.6 and a p value of ≤ 0.05. For further details and the complete list of genes, see **Supplementary Dataset 4. B.** Plot of the microarray analysis in Δ*ssr3189* and wild type under standard conditions as log_2_ fold expression of transcriptional unit (TU) TU3332, containing the genes *cmpABCD* and *slr0042*. The wild type is shown in turquoise, and Δ*ssr3189* in red. **C.** Same plot type as in panel (B) but for TU1023 with the genes *nrtABCD* and *narB*.

The genes downregulated in Δ*ssr3189* were mainly related to nitrogen uptake and metabolism (**Figure 4A**). The most down-regulated gene was *cynS* encoding cyanase (Harano et al., 1997), followed by *glnN* encoding glutamate–ammonia ligase GSIII, and the *nrtABCD* nitrate/nitrite transport system (**Figure 4C**). Another gene involved in nitrogen uptake, *amt1*, encoding ammonium permease, was also significantly downregulated, together with *nirA* and *narB* encoding nitrate reductase and ferredoxin-nitrite reductase, respectively. Both enzymes catalyze the stepwise reduction of nitrate to ammonia for the incorporation into primary metabolism in the GS/GOGAT cycle (Flores and Herrero, 1994; Flores and Herrero, 2005). These findings would be expected if there was an excess of nitrogen sensed in the cells.

Overall, the data indicated that the deletion of *ssr3189* led to a transcriptomic reorganization of several C_i_-and nitrogen-related genes.

### Intact Transcriptomic Nitrogen Starvation Response in the Δ*ssr3189* Mutant

Because of the strong effects on the transcript accumulation of C_i_-and nitrogen-related genes, we performed another transcriptome analysis, focusing on the response after 3 h and 24 h of nitrogen deprivation. This experiment revealed that the previously described transcriptomic responses to nitrogen starvation such as the induction of the *sll0783-sll0787* gene cluster (Schlebusch and Forchhammer, 2010), of *nrt, nblA* and *nir* genes (Klotz et al., 2016; Giner-Lamia et al., 2017) were clearly detected in the wild type as well as in Δ*ssr3189* (**Figure S5**). However, the magnitudes of the response differed between both strains, resulting in 31 upregulated and 106 downregulated genes in Δ*ssr3189* relative to wild type after 3 h of nitrogen starvation (**Figure 5A**). Among the upregulated genes was again *apcI* (*sll1911*) encoding an alternative linker protein (Espinoza-Corral et al., 2025). Among the downregulated genes were *cpcG2* (*sll1471*), encoding a linker protein for a photosystem I-specific antenna (Gao et al., 2016), *sll7062-7067* encoding the Cas proteins of the CRISPR2 system (Bilger et al., 2024), several genes located on plasmid pSYSM which are involved in the synthesis of synechan (Maeda et al., 2021), and, interestingly, also genes encoding regulatory proteins, *rre37* (*sll1330*) and the RNA polymerase sigma factor *sigE* (*sll1689*). Rre37 stimulates the accumulation of 2-oxoglutarate (2-OG) and glycogen under nitrogen starvation (Joseph et al., 2014), while in parallel with SigE it can activate the transcription of several sugar catabolic genes (Azuma et al., 2011). Among the targets of SigE are *gnd* encoding 6-phosphogluconate dehydrogenase and *talB* encoding transaldolase (Osanai et al., 2005; Osanai et al., 2011), while *gap1* (encoding glyceraldehyde-3-phosphate dehydrogenase) was repressed in a *rre37* disruption mutant, particularly under nitrogen-starved conditions (Azuma et al., 2011). We found all three genes repressed in the Δ*ssr3189* mutant relative to wild type (**Figure 5A**). After 24 h of nitrogen starvation, 19 genes were upregulated, and 54 genes were downregulated in Δ*ssr3189* compared to wild type (**Figure 5B**). The upregulated genes included a cassette of four not well-characterized genes (*sll1158–sll1161*), and genes involved in the carbon-concentrating mechanism (*ccmK1, ccmK2, ccmL* and *cmpC*). Relatively less expressed in Δ*ssr3189* were, again, genes on plasmid pSYSM involved in synechan synthesis, and *sll1330* encoding the response regulator Rre37.

**Figure 5.**
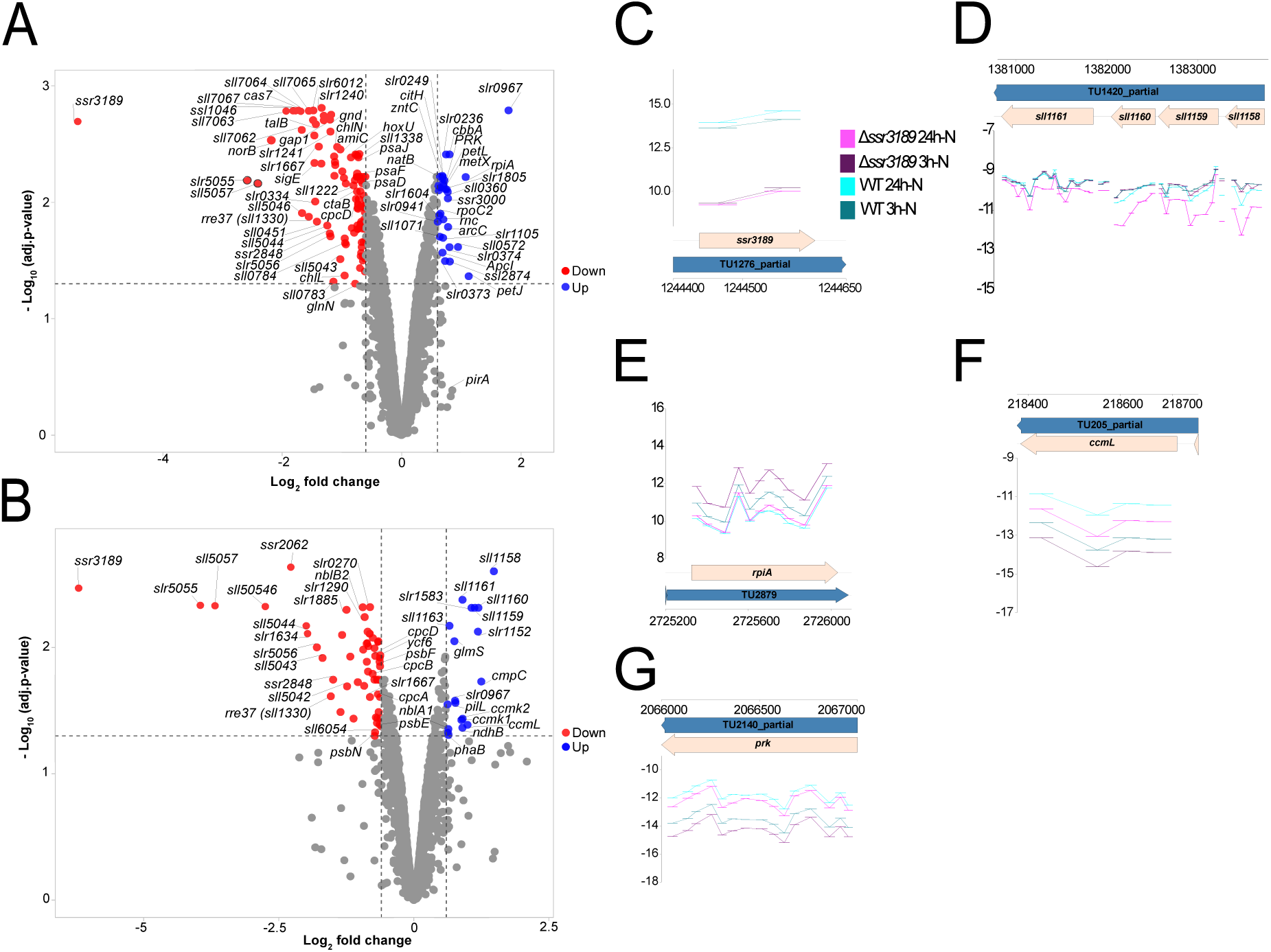
Deletion of *ssr3189* and transcriptional changes under nitrogen depletion. **A.** Microarray analysis of wild type compared to Δ*ssr3189* after 3 h of nitrogen depletion. Upregulated genes are shown in blue and downregulated genes in red. Thresholds were set at |log_2_FC| ≥ 0.6 and a p value of ≤ 0.05. For further details and the complete list of genes, see **Supplementary Dataset 4. B.** Volcano plot of microarray analysis of wild type compared to Δ*ssr3189* after 24 h of nitrogen depletion. **C.** Plot of the microarray analysis as log_2_ fold expression of the *ssr3189 gene* in Δ*ssr3189* and WT after 3 h and 24 h of nitrogen depletion. WT is shown in green (3h - N) and turquoise (24h-N), and the *ssr3189* deletion mutant is shown in purple (3h-N) and pink (24h-N). **D.** Same plot type as in panel (C), but for TU1420 with *sll1158, sll1159, sll1160,* and *sll1161*. **E.** Same plot type as in panel (C), but for gene *rpiA,* encoding ribose 5-phosphate isomerase; **F.** *ccmL,* encoding carboxysome protein L; **G.** *prk,* encoding phosphoribulokinase.

The transcriptome analysis results under standard conditions suggested an inverse correlation between the expression of genes involved in C_i_ and in nitrogen metabolism in Δ*ssr3189* in comparison to the wild type, while the analysis after 3 and 24 h of nitrogen removal showed that the nitrogen starvation response was, in principle, intact. Moreover, the differences were quantitatively relatively mild, with maximum log_2_FC differences of +2.55 (*cmpA*) and-2.07 (*cynS*, **Figure 4A**) under standard conditions and log_2_FC differences of +1.48 (*sll1158*) and-3.94 (*slr5055*) after 24 h of nitrogen starvation (**Figure 5B**).

### Metabolic Changes in the Nitrogen and Carbon Pathways

Upon *ssr3189* deletion, the phenotype (**Figure 2**), together with the observed differential expression of a specific set of genes, (**Figure 4** and **5**), pointed at effects on processes related to nitrogen and C_i_ assimilation and metabolism. To test if there was an impact on the accumulation of relevant metabolites, we performed targeted metabolomics based on liquid chromatography mass spectrometry (LC-MS/MS). We measured three biologically independent samples for wild type, Δ*ssr3189*, and Δ*ssr3189::ssr3189*×3F. In total, 31 compounds, including almost all amino acids (Cys was below the detection limit), intermediates of the arginine catabolism pathway, and several organic acids, were quantified (**Figure S6, S7**, and **Supplemental Dataset 5**). Neither the total level (**Figure S7A**), nor the individual amounts of the majority of amino acids differed significantly between wild type and Δ*ssr3189*. However, the total amount of other quantified metabolites, primarily organic acids, was significantly higher in Δ*ssr3189* than in the wild type (**Figure S7B**), caused by increased concentrations of several tricarboxylic acid (TCA) cycle metabolites, most pronounced from its oxidative branch (citrate, isocitrate, and 2-OG; **Figure 7**).

In contrast to the similar total level of amino acids, clear differences were observed for a small number of individual amino acids. In particular, glutamine was significantly (∼12 x) higher accumulated in Δ*ssr3189* than in the wild type (**Figure 6**), while it was lower (although not at wild type levels) in the complementation mutant. Significantly higher levels in Δ*ssr3189* compared to wild type were also observed for lysine (4.9 x) and arginine (3.1 x) (**Figure S6, S7C**). However, a few amino acids were also accumulated at lower levels in the Δ*ssr3189* mutant. Especially, the concentration of glutamate was significantly lower in Δ*ssr3189* compared to wild type (**Figure 6**). The discrepancy between the lower levels of glutamate and the dramatically higher levels of glutamine in Δ*ssr3189* pointed at a possibly changed glutamine synthetase activity. Therefore, the enzymatic activity of glutamine synthetase was measured, yielding 0.872 (±0.042) U/mg protein in the wild type and 1.076 (±0.117) U/mg protein in Δ*ssr3189*. Statistical testing showed that this difference in the enzymatic activity was significant (**Figure 6B**). This was a striking result, given the lowered *glnN* and (non-significantly) lowered *glnA* mRNA levels and the (non-significantly) slightly higher mRNA levels of the glutamine synthetase inactivating factors IF7 (*gifA*) and IF17 (*gifB*) in Δ*ssr3189* (**Supplemental Dataset 4**). However, Western blots showed that the glutamine synthetase amounts at protein level, were similar in Δ*ssr3189* and wild type (**Figure S8**). These results pointed at the enhanced enzymatic activity of glutamine synthetase, leading to the higher amounts of glutamine while draining the glutamate pool.

**Figure 6.**
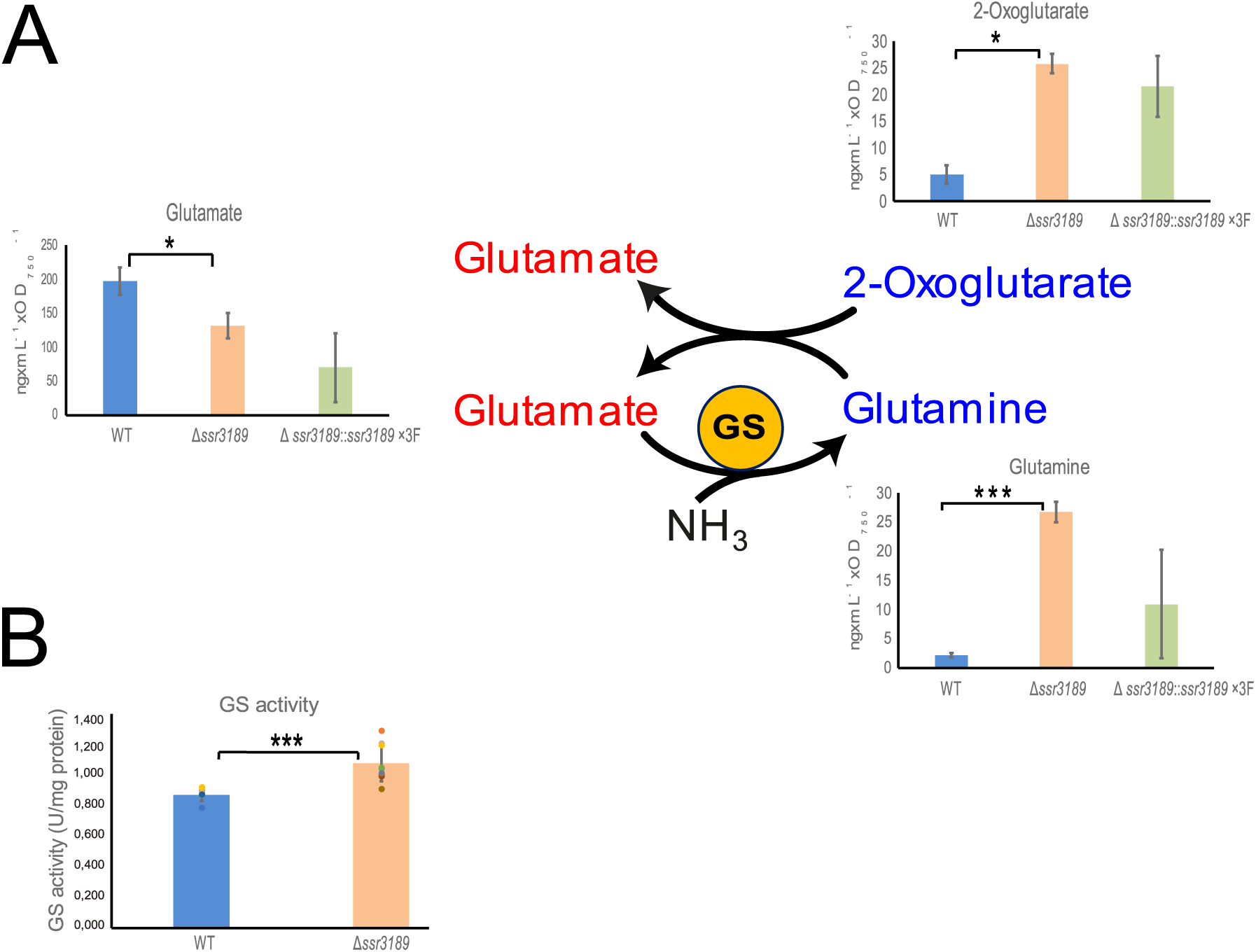
Accumulation of critical metabolites for nitrogen assimilation in *Synechocystis* 6803. **A.** Comparison of glutamine, 2-OG, and glutamine content (ng × mL^-1^ × OD_750_^-1^) in WT, deletion mutant Δ*ssr3189,* and complementation strain Δ*ssr3189*::*ssr3189*×3F. Blue labels indicate metabolites with significantly higher abundance, red labels those with significantly lower abundance in Δ*ssr3189* relative to WT. **B.** Glutamine synthetase enzymatic activity in triplicates of two parallel wild type cultures (n=6) and three parallel Δ*ssr3189* mutant lines (n=9). The activity was significantly higher in the mutant. Significance was calculated with two-sample *t* test with unequal variance (Welch’s *t* test, *; *P* < 0.05, **; *P* < 0.01, ***; *P* < 0.001), see **Supplementary Table S1** for the individual numbers.

**Figure 7.**
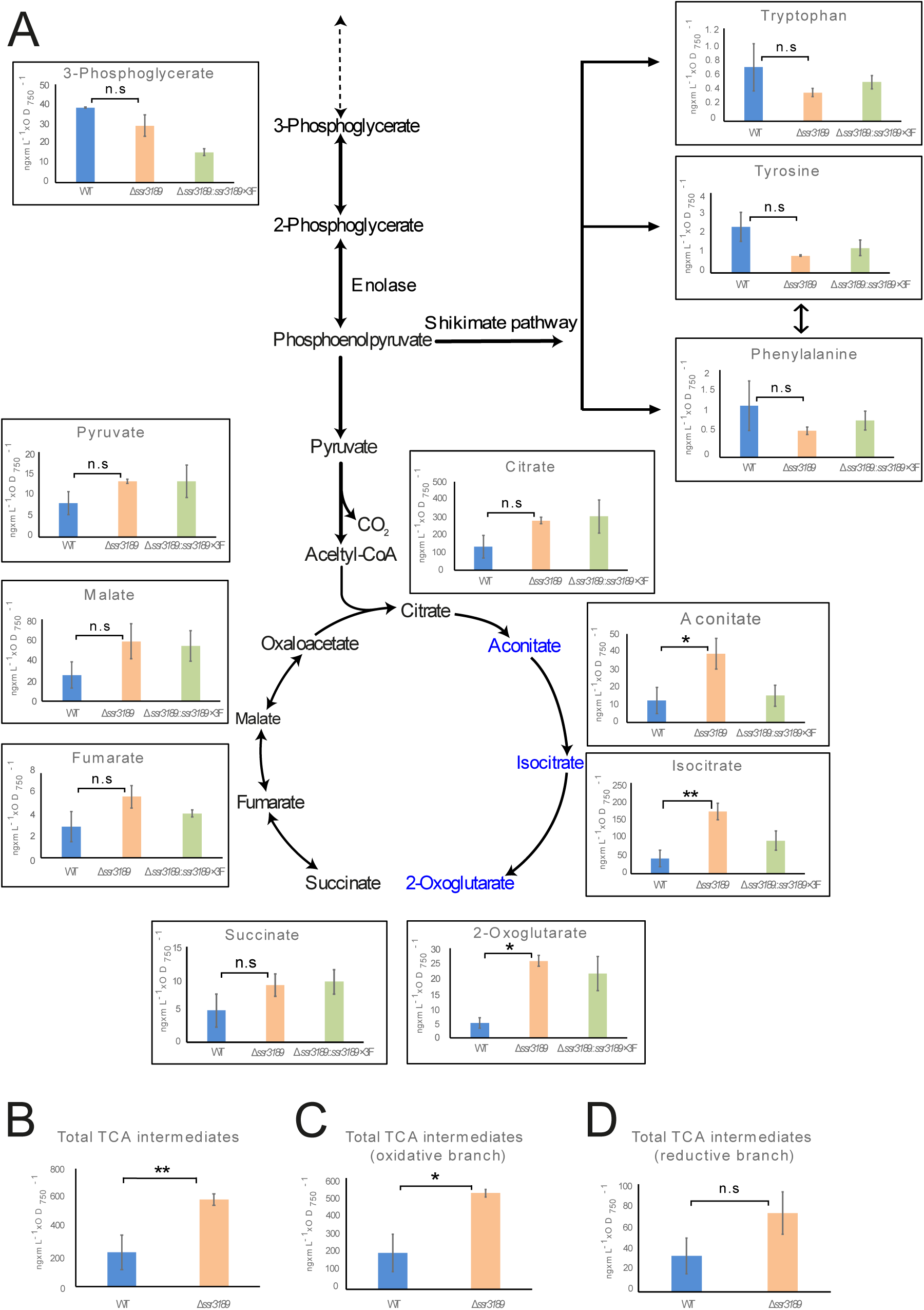
Accumulation of TCA cycle metabolites. **A.** Comparison of metabolites content (ng × mL^-1^ × OD_750_^-1^) in the wild type (WT), deletion mutant Δ*ssr3189,* and complementation strain Δ*ssr3189*::*ssr3189*×3F under standard growth conditions (Acetyl CoA, Acetyl coenzyme A). **B.** Total TCA intermediates, **C.** Total TCA intermediates (oxidative branch). **D.** Total TCA intermediates (reductive branch). Blue labels indicate metabolites with significantly higher abundance in Δ*ssr3189* relative to WT. Significance was calculated with two-sample *t* test with unequal variance (Welch’s *t* test, n.s; not significant, *; *P* < 0.05, **; *P* < 0.01, ***; *P* < 0.001).

Interestingly, the total amounts of other quantified metabolites, primarily organic acids, were significantly higher in cells of the Δ*ssr3189* mutant compared to wild type (**Figure S7B**). Among the most relevant metabolites, we measured significantly (2.8 x) more 2-OG in the mutant (**Figure 6A**), indicating that the amounts of 2-OG were not limiting for glutamate formation. In addition, as the key nitrogen status-sensing molecule (reviewed by Forchhammer and Selim (2020)), the higher 2-OG level should signal nitrogen starvation in the Δ*ssr3189* mutant.

Our measurements furthermore showed that the higher levels of organic acids included several metabolites of the TCA cycle, most pronounced for aconitate and citrate (**Figure 7**). Of further interest was 3-PGA, which is connected to the formation of aromatic amino acids via the shikimate pathway. We noticed that 3-PGA, the aromatic amino acids phenylalanine, tryptophan and tyrosine (**Figure 7**), as well as histidine (**Figure S7C**) had lower (but not significantly lower) concentrations in Δ*ssr3189* than in the wild type.

Collectively, the metabolome data clearly supported that the protein encoded by *ssr3189* plays an important role in the regulation of C/N homeostasis in *Synechocystis* 6803.

### Co-immunoprecipitation Reveals Enolase, the CRISPR Endoribonuclease Cas6-1, and Ribosomal Proteins as Interaction Partners

Co-IP experiments were performed to gain further insight into the regulatory-metabolic network in which *ssr3189* is involved. The elution samples showed in Western blots a strong signal of a 15 kDa protein expressed from P*_petE_*-*ssr3189*×3F (**Figure S9A**). Further bands corresponding to higher molecular masses, most pronounced for a signal at 35 kDa were detected, too, as seen in the Western blot of the complementation strain (**Figure 2B**). To test the likelihood of complex formation, a higher amount of protein was loaded, and the samples were kept in native or denatured conditions. The membrane showed several bands, some of which were only present in the native conditions, indicating interactions with further proteins or multimeric complexes (**Figure S9B**).

The analysis of the co-IP elution samples by mass spectrometry identified 88 enriched proteins together with Ssr3189 (**Figure 8, Supplemental Dataset 6**). The 78 most enriched proteins (log_2_FC ≥3) are given in **Table 1**. Together with the bait protein, enolase, a key enzyme in the central carbon metabolism of *Synechocystis* 6803 (Knowles and Plaxton, 2003), and the CRISPR1 endoribonuclease Cas6-1 (Reimann et al., 2017) were the most enriched proteins (**Table 1**). Among the other interactors were all 56 ribosomal proteins, as well as proteins associated with the ribosome or the translation process, such as the translation initiation factor InfC, the tmRNA-binding protein SsrA required for releasing stalled ribosomes, the ribosome maturation factor RimP, and Sll1830, which is predicted to be structurally similar to a subunit of the peptide chain release factor. Only a few metabolic enzymes were among the enriched proteins, in addition to enolase these were arginine decarboxylase 1 and 2, porphobilinogen synthase HemB (Delta-aminolevulinic acid dehydratase), pyruvate dehydrogenase (E1) component catalyzing the rate-limiting first step of the pyruvate dehydrogenase complex to produce the TCA cycle precursor acetyl-CoA, and AcnB catalyzing the isomerization of citrate to isocitrate via cis-aconitate, a vital step in the TCA cycle.

**Figure 8.**
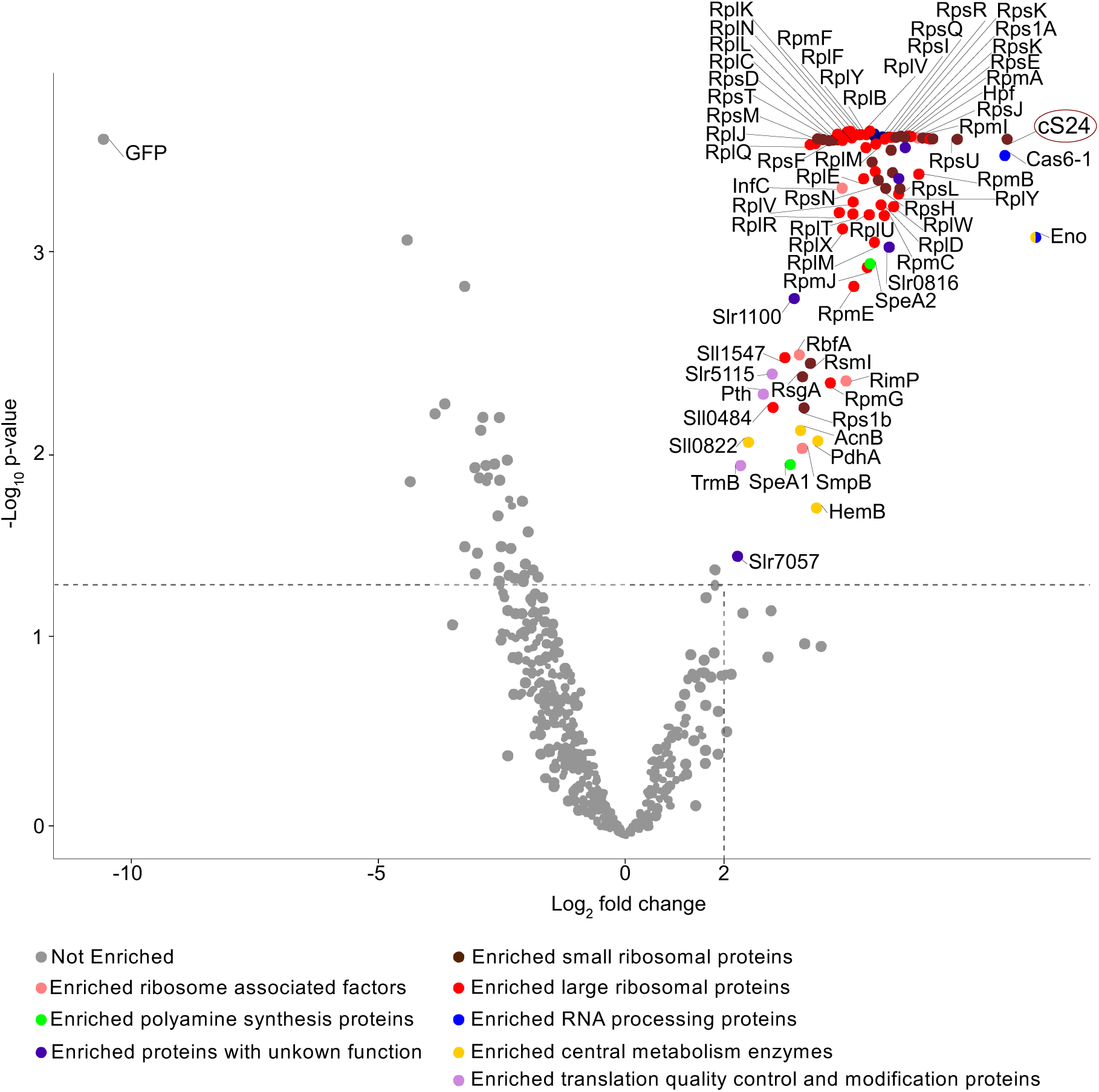
Co-immunoprecipitation analysis for the identification of proteins co-enriched with Ssr3189. Volcano plot of detected proteins in the P*_petE_*-*ssr3189*×3F elution fraction compared with the P*_petE_*-*s p*×3F control based on a Limma-moderated t-test with adjustment for multiple testing correction. The thresholds for p value and log_2_ fold change were set to 0.05 and 2, respectively. For the complete list of detected proteins, see **Supplemental Dataset 6**.

**Table 1.** Proteins enriched together with Ssr3189 by co-IP. The entries are ranked according to the enrichment fold change of triplicate LFQ values (log_2_FC). Only proteins with a log_2_FC ≥ 3 are shown. Statistical significance of enriched proteins is represented by-log_10_ P value ≥ 1.3. Abbreviations: 2-PG, 2-phosphoglycerate; LSU, large subunit; PEP, phosphoenolpyruvate; SSU, small subunit; r-protein, ribosomal protein. The bait protein was identified based on a unique peptide (Ssr3189-DYK-FLAG). For visualization, see **Figure 8**, for the complete dataset, see **Supplemental Dataset 6**.

| Rank | UniProt ID | Protein | Description | $\log_2FC$ | $-\log_{10} P$<br>value |
| --- | --- | --- | --- | --- | --- |
| 1 | P77972 | Enolase | converting 2-PG into PEP | 8.27 | 3.18 |
| 2 | P73656 | Ssr3189 | bait | 7.68 | 3.71 |
| 3 | Q6ZEI4 | Cas6-1 | CRISPR1 endoribonuclease | 7.64 | 3.63 |
| 4 | P48949 | RpsU | SSU r-protein S21 | 6.67 | 3.71 |
| 5 | P73314 | RpsC | SSU r-protein | 6.09 | 3.71 |
| 6 | P74071 | RpsB | SSU r-protein | 6.05 | 3.71 |
| 7 | P73316 | RpsS | SSU r-protein | 6.03 | 3.71 |
| 8 | P48959 | RpmI | SSU r-protein | 6.02 | 3.71 |
| 9 | P74226 | RpsJ | SSU r-protein | 5.92 | 3.71 |
| 10 | P73293 | RpsI | SSU r-protein | 5.89 | 3.71 |
| 11 | P72851 | RpmB | LSU r-protein | 5.89 | 3.53 |
| 12 | P74518 | Hpf | ribosome hibernation promotion factor | 5.88 | 3.71 |
| 13 | P42352 | RplI | LSU r-protein | 5.83 | 3.71 |
| 14 | P74267 | RpmA | LSU r-protein | 5.83 | 3.71 |
| 15 | P73304 | RpsE | SSU r-protein | 5.76 | 3.71 |
| 16 | P73298 | RpsK | SSU r-protein | 5.75 | 3.71 |
| 17 | P73530 | Rps1A | SSU r-protein | 5.73 | 3.71 |
| 18 | P73120 | Sll1830 | peptide chain release factor | 5.65 | 3.71 |
| 19 | Q55555 | Sll0175 | NAD(P)/FAD-dependent oxidoreductase | 5.62 | 3.67 |
| 20 | P48946 | RpsR | SSU r-protein | 5.60 | 3.71 |
| 21 | P73311 | RpsQ | SSU r-protein | 5.56 | 3.71 |
| 22 | P74442 | Slr0143 | WD repeat-containing protein | 5.54 | 3.71 |
| 23 | P74230 | RpsL | SSU r-protein | 5.50 | 3.45 |
| 24 | P73289 | RplY | LSU r-protein | 5.50 | 3.42 |
| 25 | P73317 | RplB | LSU r-protein | 5.49 | 3.71 |
| 26 | Q6YRW9;<br>Q6ZEM4 | Ssl6035;<br>Ssr5106 | unknown | 5.49 | 3.50 |
| 27 | P73306 | RplF | LSU r-protein | 5.48 | 3.71 |
| 28 | P36237 | RplK | LSU r-protein | 5.43 | 3.71 |
| 29 | P73313 | RplP | LSU r-protein | 5.41 | 3.71 |
| 30 | P72866 | RpsO | SSU r-protein | 5.36 | 3.54 |
| 31 | P73318 | RplW | LSU r-protein | 5.36 | 3.35 |
| 32 | P23349 | RplL | LSU r-protein | 5.35 | 3.71 |
| 33 | P74410 | RpsP | SSU r-protein | 5.33 | 3.66 |
| 34 | P74049 | Slr0816 | LSU accumulation protein YceD | 5.29 | 3.13 |
| 35 | P73636 | RpsF | SSU r-protein | 5.26 | 3.71 |
| 36 | P74229 | RpsG | SSU r-protein | 5.26 | 3.71 |
| 37 | P73312 | RpmC | LSU r-protein | 5.24 | 3.31 |
| 38 | P73307 | RpsH | SSU r-protein | 5.22 | 3.45 |
| 39 | P73310 | RplN | LSU r-protein | 5.21 | 3.71 |
| 40 | P48939 | RpsD | SSU r-protein | 5.21 | 3.71 |
| 41 | P73320 | RplC | LSU r-protein | 5.18 | 3.71 |
| 42 | P73014 | RpmF | LSU r-protein | 5.16 | 3.71 |
| 43 | P73299 | RpsM | SSU r-protein | 5.16 | 3.71 |
| 44 | P73319 | RplD | LSU r-protein | 5.13 | 3.36 |
| 45 | P73296 | RplQ | LSU r-protein | 5.11 | 3.71 |
| 46 | P73303 | RplO | LSU r-protein | 5.07 | 3.49 |
| 47 | P73336 | RpsT | SSU r-protein | 5.03 | 3.71 |
| 48 | P23350 | RplJ | LSU r-protein | 5.01 | 3.71 |
| 49 | P48944 | RpsN | SSU r-protein | 5.01 | 3.54 |
| 50 | P73294 | RplM | LSU r-protein | 4.99 | 3.16 |
| 51 | P36239 | RplS | LSU r-protein | 4.95 | 3.68 |
| 52 | P36236 | RplA | LSU r-protein | 4.95 | 3.59 |
| 53 | P74266 | RplU | LSU r-protein | 4.90 | 3.28 |
| 54 | P72587 | SpeA2 | arginine decarboxylase 2 | 4.88 | 3.04 |
| 55 | P73300 | RpmJ | LSU r-protein | 4.85 | 3.03 |
| 56 | Q55385 | Slr0923 | SSU r-protein | 4.82 | 3.67 |
| 57 | P73308 | RplE | LSU r-protein | 4.78 | 3.50 |
| 58 | P73292 | RpmE | LSU r-protein | 4.58 | 2.92 |
| 59 | P73315 | RplV | LSU r-protein | 4.56 | 3.37 |
| 60 | P48957 | RplT | LSU r-protein | 4.56 | 3.31 |
| 61 | P72874 | InfC | Translation initiation factor IF-3 | 4.55 | 3.44 |
| 62 | P72687 | RimP | Ribosome maturation factor | 4.55 | 2.38 |
| 63 | P73309 | RplX | LSU r-protein | 4.37 | 3.22 |
| 64 | P73305 | RplR | LSU r-protein | 4.30 | 3.31 |
| 65 | P48958 | RpmG | LSU r-protein | 4.10 | 2.38 |
| 66 | P77969 | HemB | delta-aminolevulinic acid dehydratase | 3.97 | 1.72 |
| 67 | P74490 | PdhA | pyruvate dehydrogenase E1 subunit alpha | 3.86 | 2.08 |
| 68 | P74355 | SmpB | SsrA-binding protein | 3.72 | 2.04 |
| 69 | P74038 | RsmI | LSU r-protein | 3.67 | 2.51 |
| 70 | P74142 | Rps1b | SSU r-protein | 3.57 | 2.27 |
| 71 | Q55625 | RbfA | ribosome-binding factor A | 3.56 | 2.53 |
| 72 | P52640 | RsgA | SSU biogenesis GTPase RsgA | 3.54 | 2.43 |
| 73 | P74582 | AcnB | aconitase | 3.50 | 2.15 |
| 74 | P72744 | Slr1100 | ParE-like toxin | 3.40 | 2.84 |
| 75 | P74662 | Sll1547 | putative pre-16S rRNA nuclease | 3.36 | 2.51 |
| 76 | P74576 | SpeA1 | arginine decarboxylase 1 | 3.29 | 1.96 |

Because enolase was so highly enriched in the co-IP with Ssr3189 (**Figure 8**) and connects 3-PGA with the formation of aromatic amino acids via the shikimate pathway (**Figure 7**), we tested its enzymatic activity in crude extracts of Δ*ssr3189* and the wild type. However, the enzymatic activity of enolase was not significantly different in extracts of the Δ*ssr3189* mutant compared to wild type (**Figure S10**).

The observation that all ribosomal proteins were enriched, together with several enzymes, among them enolase, an enzyme which is forming the backbone of bacterial degradosomes (Zhang et al., 2022), was consistent with the role of the *ssr3189* gene product as ribosomal protein S24 as previously suggested (Pearce et al., 2026), and indicated a connection to metabolic regulation.

### Ssr3189 Is an RNA-binding protein

The previously found behavior of Ssr3189 in GradR analyses (Hemm et al., 2025b) raised the question if Ssr3189 is an RNA-binding protein. Analyzing these data more closely, we found Ssr3189 in the not RNase-treated GradR gradient in the bottom fractions, co-fractionating with ribosomes (**Figure 9A**). Upon RNase treatment, the majority of Ssr3189 shifted to fractions 8 to 12, indicating it had been part of an RNA-containing complex that was dismantled when the RNA was degraded, a minor fraction even showed up in the lightest (top) fraction, likely representing monomeric protein (**Figure 9A**). Moreover, recent structural analyses of the *Synechocystis* 6803 ribosome reported the proximity of this protein to the 16S ribosomal RNA (Pearce et al., 2026). These observations suggested, but did not prove that Ssr3189 was directly binding RNA. Therefore, we used isotope labeling by PNK (Brenes-Álvarez et al., 2025) to test for RNA-binding directly. We used the P*_petE_*-*ssr3189*x3F strain expressing the 3xFLAG epitope-tagged protein under control of the copper-inducible P*_petE_* promoter (**Figure S3A**). In parallel, we used a strain expressing another nucleic acid-binding protein, the SyCrp1 transcription factor (gene *sll1371*) as a negative control (Hemm et al., 2025b). After UV-treatment to covalently crosslink the protein to the possibly associated RNA, the complexes were purified via magnetic beads binding to the FLAG epitope tag under stringent conditions to disrupt protein-protein interactions. Then, the RNA was partially degraded by benzonase to select for nuclease-protected RNA covalently bound to the protein. The remaining RNA fragments were ^32^P-isotope labeled by PNK, followed by electrophoretic separation on SDS-PAA gels and transfer to nitrocellulose membrane. This assay yielded a clear signal for Ssr3189 indicating RNA binding, while no signal was obtained for SyCrp1 (**Figure 9B**). Western blotting and incubation with an anti-FLAG antiserum validated the presence of both tagged proteins irrespectively of the UV-treatment (**Figure 9C**).

**Figure 9.**
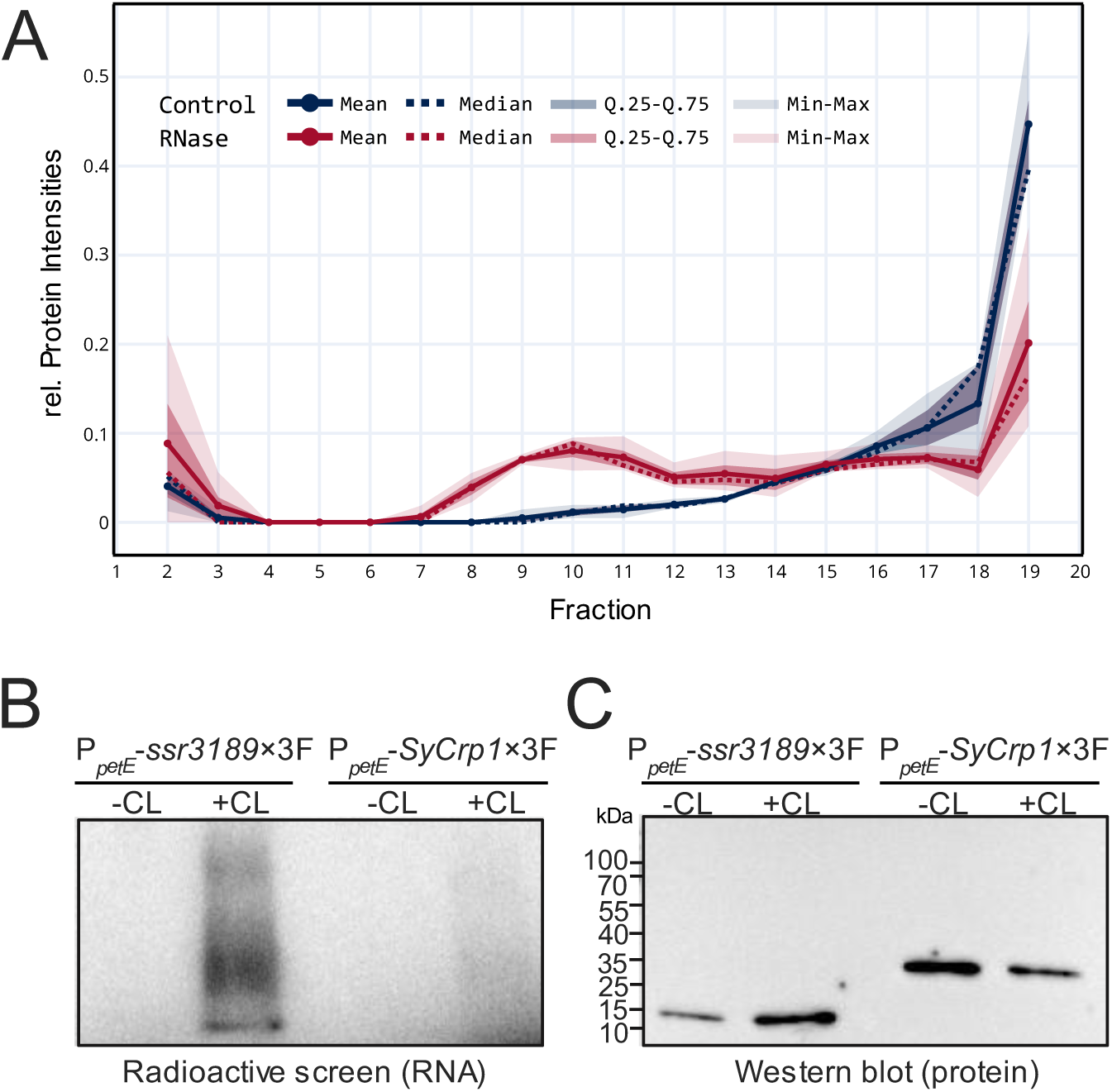
Ssr3189 is an RNA-binding protein. **A.** Fractionation analysis by sucrose gradient ultracentrifugation in the presence (rubin) or absence (dark blue) of RNase (Hemm et al., 2025b). The mean and median values are plotted from measured triplicate cultures. **B.** Radioactive screen of the Western blot membrane of the PNK assay of Ssr3189 (expressed from P*_petE_*-*ssr3189*×3F and SyCRP1 (expressed from P*_petE_*-*SyCrp1*×3F, negative control). **C.** Western blot analysis of the PNK assay (-CL: not UV-crosslinked; + CL, UV-crosslinked).

We conclude that these results validated Ssr3189 as an RNA-binding protein.

## DISCUSSION

### Ssr3189 as an RNA-Binding Ribosomal Protein

Previous GradR analyses in *Nostoc* sp. PCC 7120 identified the homolog of Ssr3189, Asl3888 as an RNA-binding protein (Brenes-Álvarez et al., 2025) which led to the definition of the RNA-binding Pfam domain PF30515 (https://www.ebi.ac.uk/interpro/entry/pfam/PF30515). Structural analysis of the *Synechocystis* 6803 ribosome showed Ssr3189 named as bS24 associated to the ribosome, close to the 16S ribosomal RNA, but did not show direct interaction (Pearce et al., 2026). Our data validate Ssr3189 as an RNA-binding protein. Several ribosomal proteins are known in bacterial model organisms to be RNA-binding. These proteins often bind rRNA forming the ribosomal ribonucleoprotein complex, but some bind also their own mRNAs interfering with translation. Most mRNAs encoding ribosomal proteins are multicistronic, therefore, these RNA-binding ribosomal proteins are central elements of a regulatory feedback mechanism (Nomura et al., 1980; Yates and Nomura, 1981). These processes are not studied in cyanobacteria, but similar mechanisms can be expected. All the more it is intriguing that Ssr3189, a protein closely associated with the cyanobacterial ribosome, is RNA-binding and encoded by a gene that does not belong to one of the classical ribosomal protein operons. The presence of *ssr3189* homologs in several distinct lineages of alga (and also *Cercozoa* and haptophytes) strengthens the argument that Ssr3189 is an important protein that originated in cyanobacteria, was retained in various algae after endosymbiosis, but was lost in plant chloroplasts. Consistent with its presence in most cyanobacteria and the chloroplasts of several algae, this protein family might be called cS24 (cyanobacterial/chloroplast ribosomal protein S24).

Together with Rbp3 (Hemm et al., 2025a) and YlxR/RnpM (Hemm et al., 2024), cS24 belongs to a very small group of verified RNA-binding proteins in *Synechocystis* 6803. As a ribosomal protein, cS24 was found close to the anti-Shine-Dalgarno motif within the 16S rRNA, which prompted the suggestion that it may contribute to the recognition of the Shine-Dalgarno motif during the formation of the ribosomal initiation complex, perhaps similar to the RpsU/S21 protein (Pearce et al., 2026). Consistent with these ideas, we noticed that the RpsU/S21 protein was the most highly enriched ribosomal protein in our co-IP with the Flag-tagged cS24 protein (**Figure 8**, **Table 1**).

However, most genes in *Synechocystis* 6803 lack a classical Shine-Dalgarno motif (Hadjeras et al., 2025). We previously also noticed a 3 or 4 nt offset in the mapping of ribosomal footprints stalled on start and stop codons in *Synechocystis* 6803 compared to *E. coli* and *Campylobacter jejuni* (Hadjeras et al., 2025), pointing at further differences in the architecture of the ribosome between cyanobacteria and better studied other bacteria.

### The Ssr3189 Interactome

Small proteins frequently function as regulators, modulating the activity of enzymes or large protein complexes through binding to them. For a few small proteins, the protein interactome was characterized using co-IP analyses, such as NblD, AtpΘ or NirP1 (Krauspe et al., 2022; Song et al., 2022, Kraus et al., 2024). The same method was used in the present study. While the co-enrichment of all ribosomal proteins with cS24 is consistent with its recent identification as ribosomal protein (Pearce et al., 2026), we also found strongly enriched other proteins, especially enolase, and Cas6-1 (**Figure 8**, **Table 1**). This points at the possibility that cS24 is involved in additional interactions, possibly including degradosomes, which are multienzyme RNA-degrading complexes in bacteria (Carpousis et al., 2022), and in which this enzyme is a central component in many bacteria (Zhang et al., 2022). However, none of the degradosome ribonucleases were co-enriched making this possibility less likely. However, the co-enrichment of the CRISPR1 endoribonuclease Cas6-1 was pointing in the direction of RNA processing. This endoribonuclease is the critical factor in the maturation of CRISPR RNAs (crRNAs) in the CRISPR1 system, one of three CRISPR-Cas systems in *Synechocystis* 6803 (Scholz et al., 2013). *In vitro* studies revealed that Cas6-1 can also process the crRNAs of the CRISPR2 system (Reimann et al., 2017). We observed a strong downregulation of mRNAs for the CRISPR2 Cas proteins (**Figure 4A, 5A**) in the Δ*ssr3189* strain. Therefore, cS24 could be involved in mediating cross-talk between the CRISPR1 and CRISPR2 systems. Alternatively, cS24 may sequester Cas6-1 for targeting truncated or damaged crRNAs to the degradosome. We notice that interactions with ribosomes, possibly degradosomes, and the CRISPR machinery would be consistent with the capability of cS24 to bind RNA as these are all RNA-protein complexes.

Although with somewhat lower enrichment factors, the cS24 co-IP yielded also several additional metabolic enzymes (**Table 1**). The possibility that these interactions impact the respective metabolic pathways is an intriguing possibility.

### Absence of *ssr3189* Impacts the C/N Metabolism

We observed multiple alterations in the accumulation of amino acids and organic acids in the Δ*ssr3189* mutant compared to wild type. Most pronounced was the overaccumulation of glutamine (**Figure 6A**). This effect can be related to two facts. First, the production of its precursors, i.e., TCA intermediates in its oxidative branch. In cyanobacteria, the TCA cycle is considered to be largely open, because the 2-OG dehydrogenase complex is missing. While shunts potentially closing the cyanobacterial TCA cycle have been reported, the flux through these shunts appears to be rather low (Zhang and Bryant, 2011; Xiong et al., 2014). Therefore, the oxidative TCA branch initiated by aconitase leads in cyanobacteria mainly to the production of 2-OG as the precursor of ammonia assimilation by glutamine synthetase and glutamate synthase (GOGAT). We measured a higher accumulation of citrate, aconitate and 2-OG in Δ*ssr3189*, providing plenty of carbon skeletons for the glutamine synthetase-GOGAT cycle.

Second, we measured a significantly higher enzymatic activity of glutamine synthetase (**Figure 6B**), which caused a depletion in its immediate substrate, glutamate. Therefore, the replenishment of Glu through GOGAT is the likely bottleneck, while the overaccumulation of glutamine indicates that it could not efficiently be utilized by the cells.

Likely connected is our observation of elevated levels for several other amino acids, in particular lysine and arginine. However, aromatic amino acids were less accumulated. These amino acids share their synthesis via the shikimate pathway, which starts with the enzyme enolase, which was also highly co-enriched with cS24 in the co-IP experiment. Enolase is not only a component of the degradosome in many bacteria, but also a key enzyme in the central carbon metabolism of *Synechocystis* 6803 (Knowles and Plaxton, 2003). Moreover, the experimental repression of enolase limited the synthesis of aromatic amino acids in plant chloroplasts (Voll et al., 2009). We therefore tested the enolase enzymatic activity in crude extracts of Δ*ssr3189* and the wild type. However, the enolase activity was not significantly different (**Figure S10**), pointing at the involvement of other, unknown factors in causing this effect.

But what led to the higher accumulation of organic acids in the Δ*ssr3189* mutant in the first case? One factor could be direct interaction between cS24 and relevant enzymes. We noticed that three of the co-enriched enzymes occupy central positions in the respective pathways. In addition to enolase, these were pyruvate dehydrogenase E1 subunit alpha, the core catalytic component of the pyruvate dehydrogenase (PDH) complex that connects glycolysis to the TCA cycle (Wang et al., 2022), and aconitase catalyzing the isomerization of citrate to isocitrate, a vital step in the TCA cycle. If cS24 regulates the activities of some enzymes directly, it likely does so through its C-terminal half. In contrast to the very conserved and strongly electropositive N-terminal section, this segment is less conserved and predicted as a disordered region (**Figure 1C** and **2B**), making it a candidate for such interactions. We also noticed that complementation in strain Δ*ssr3189*::*ssr3189*×3F worked well for some phenotypical effects, but failed for others, possibly due to the C-terminal FLAG tag hampering some, but not all interactions. Therefore, the possible effects of cS24 on these enzymes are an interesting topic for further research.

However, another likely relevant factor contributing to the accumulation of organic acids in the Δ*ssr3189* mutant is our observation of the enhanced expression of several genes directly involved in the uptake of C_i_ from the environment such as the *cmpABCD* operon (**Figure 4** and **5**). At the same time, we observed a downregulation in the expression of uptake systems for nitrogen, such as *amt1*, or the *nrtABCD* operon (**Figure 4**). NtcA is the transcription factor which controls several of these operons (Giner-Lamia et al., 2017) and which directly responds to the concentration of 2-OG (Zhao et al., 2010). Hence, the signs for intracellular nitrogen starvation observed by the pigmentation phenotype and the altered gene expression in the mutant under standard growth conditions could be sensed via the elevated 2-OG amounts in the mutant cells. This prompted us to expose the Δ*ssr3189* mutant for 3 or 24 h of nitrogen starvation to test the NtcA-mediated response. Despite some quantitative differences, the results showed that the nitrogen starvation response was qualitatively intact in the Δ*ssr3189* mutant (**Figure S5**).

### A Connection between Protein Synthesis, Primary Metabolism and RNA Degradation?

Our data connect cS24 to protein synthesis on ribosomes, but also to primary metabolism and, possibly, RNA degradation (**Figure 10**). The link to RNA degradation is speculative at the present time, because in addition to enolase, Cas6-1 was the only co-enriched ribonuclease, while classical degradosome enzymes such as RNase E or RNase J were not enriched. However, the alterations in metabolite levels in the Δ*ssr3189* mutant were remarkable. This points at a scenario where the cells became flooded with C_i_ that could only partially be metabolized further due to the lack of assimilated nitrogen. Changes in the accumulation and translation of certain mRNAs could possibly explain these findings. Therefore, it is noteworthy that, besides their canonical function in translation, several r-proteins display moonlighting activities. For example, bS1 functions also in translational control, transcription, and RNA decay (for reviews, see (Hajnsdorf and Boni, 2012; Aseev et al., 2024)). The protein bS21-2, one of the three bS21 homologs in the human pathogen *Francisella tularensis*, facilitates the preferential translation of virulence genes based on the recognition of a six-nt sequence motif in their 5′UTRs (Trautmann et al., 2023). Intriguingly, RpsU/S21was also the most highly enriched ribosomal protein in our cS24 co-IP, and both proteins were found spatially close in the structural analysis of the *Synechocystis* 6803 ribosome (Pearce et al., 2026). Consequently, cS24 could facilitate the preferred translation of a subset of mRNAs, either directly, or by mediating the interaction with regulatory small RNAs. The identification of the molecular details of its mode of action is an exciting topic for future research.

**Figure 10.**
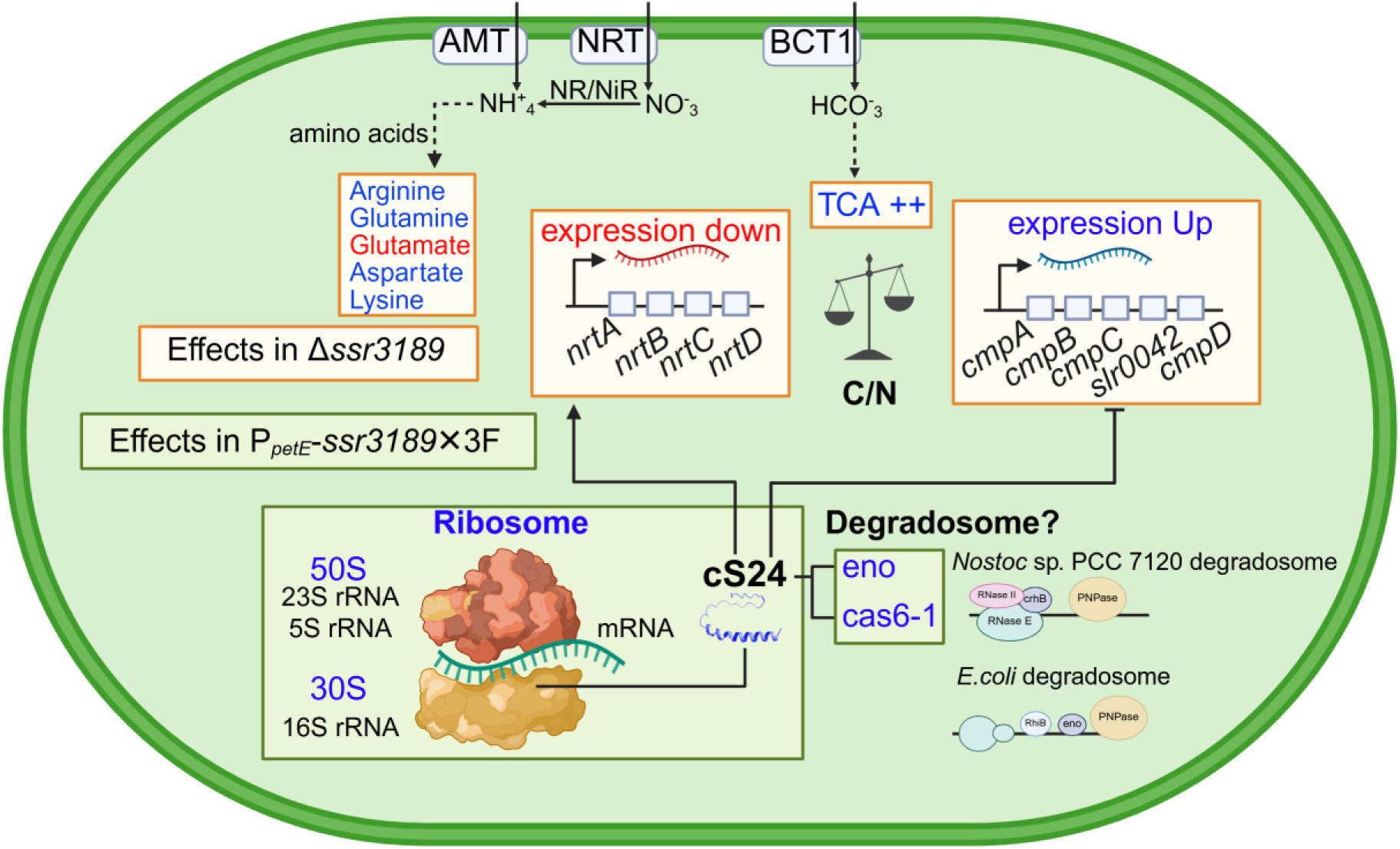
Overview of functions of cS24. Deletion of the gene *ssr3189* encoding cS24 led to changes in the transcript levels of nitrogen and carbon transporters (Figure 4), as well as metabolic changes in amino acid and organic acid levels (Figure 6, 7**, and Figure S6, S7**). Co-IP analysis revealed ribosomal proteins, enolase, and Cas6-1 as the most co-enriched proteins (Figure 8), consistent with functions related to the ribosome, carbon metabolism, as well as a possible involvement in the RNA degradation complex, the degradosome (schematically depicted for the better characterized constituents in the cyanobacterium *Nostoc* sp. PCC 7120 and in *E. coli*). The figure was made with elements from Biorender.com.

## MATERIALS AND METHODS

### Cultivation Conditions and Spectrometric Measurements

*Synechocystis* 6803 cultures were grown in BG11 or in copper-free BG11 (Rippka et al., 1979) supplemented with 20 mM TES pH 7.5 under continuous illumination with white light at 30 – 50 µmol photons m^-2^ s^-1^ at 30 °C and constant shaking. Mutant strains containing pVZ322-derived plasmids were cultivated in the presence of 50 µg/mL kanamycin, while Δ*ssr3189* was maintained at 20 µg/mL streptomycin. The cultures for overexpression of Ssr3189 using the P*_petE_* promoter for pull-down experiments and MS measurements were grown either in BG11 or under CellDeg® conditions supplemented with 2 µM CuSO_4_ as previously reported (Lippi et al., 2018; Migur et al., 2021). Cultures in growth experiments and phenotypic assays were grown in biological triplicates in BG11 medium copper-free medium and without antibiotics to prevent any possible side effects.

Whole-cell absorption spectra were measured using a Specord® 250 Plus (Analytik Jena) spectrophotometer at room temperature and were normalized to 750 nm. Cultures were measured in triplicate. Cultures at an OD_750_ > 1 were diluted with 1× BG11 prior to taking the absorption spectra.

### Generation of Mutant and Overexpression Lines

*Synechocystis* 6803 PCC-M (Trautmann et al., 2012) was used as the wild type and background for the generation of mutants. Knockout mutants were generated by replacing the *ssr3189* coding sequence with a streptomycin resistance cassette (*aadA*) (Hollingshead and Vapnek, 1985) by homologous recombination and using pUC19 as a vector for subcloning. The construct for gene replacement was generated by AQUA cloning (Beyer et al., 2015). Details of primer sequences and plasmids are provided in **Supplementary Tables S2 and S3**. Segregation of *ssr3189* and Δ*ssr3189/aadA* alleles was checked by colony PCR using the primers P_AK13 and P_AK14 (**Supplementary Figure S3B**).

To complement the knockout and generate an overexpressor and complementation strain, *ssr3189* was inserted into the self-replicating pVZ322 plasmid under control of the Cu^2+^-inducible P*_petE_* promoter or under control of its native promoter (P*_ssr3189_*). The inserts for the overexpression *of ssr3189* were assembled into pVZ322 predigested by *Xba*I and *Pst*I for 16 h at 37 °C by AQUA cloning. The constructs were checked using the primers P_AK22 and P_AK23 (**Supplementary Figure S3C**). The sequence-verified plasmids were introduced into *Synechocystis* 6803 by electroporation (Ludwig et al., 2008). The expression of *ssr3189* was checked by Western hybridization using ANTI-FLAG antisera.

### RNA Extraction, Northern Blot Verification, PNK Assay and Microarray Analysis

For transcript verification, wild type, Δ*ssr3189* and complementation strain were cultivated in Erlenmeyer flasks in BG11 media at 30 °C, under continuous illumination of 50 µmol photons m^-2^ s^-1^ with constant shaking. Samples were taken in exponential, stationary phase and 3, 6 and 24 h after nitrogen deprivation. For nitrogen deprivation experiments, exponentially growing cells were harvested by centrifugation, washed three times with nitrogen-free BG11 medium to remove residual nitrate, and resuspended in nitrogen-free BG11 medium lacking sodium nitrate (NaNO_3_) while containing 0.38 mM sodium carbonate (Na_2_CO_3_). Cultures were subsequently incubated under the same growth conditions and samples were collected at the indicated time points for RNA extraction and microarray analysis. Cells were collected by filtration through hydrophilic polyethersulfone filters (Pall Supor®−800, 0.8 μm), transferred into PGTX buffer, and instantly frozen in liquid N_2_. RNA was prepared from the frozen filter membranes as previously described (Kraus et al., 2024). The *ssr3189* transcript accumulation was analyzed by Northern hybridization using single-stranded radioactively labeled RNA probes transcribed *in vitro* from PCR-generated templates using primers P_AK24 and P_AK25 (**Supplementary Table S2**). Isotope-labeled single-stranded RNA probes were generated using [α-^32^P]-UTP and the Maxiscript T7 *In vitro* transcription kit (Thermo Fisher Scientific). Hybridizations were performed in 50% deionized formamide, 7% SDS, 250 mM NaCl, and 120 mM Na_2_HPO_4_/NaH_2_PO_4_ pH 7.2 overnight at 62 °C. The membranes were washed in buffer 1 (2× SSC (3 M NaCl,0.3 M sodium citrate, pH 7.0), 1% SDS), buffer 2 (1× SSC, 0.5% SDS) and buffer 3 (0.1× SSC, 0.1% SDS) each for 10 min. All wash steps were performed at 57 °C. Signals were detected with a Laser Scanner Typhoon FLA 9500 (GE Healthcare). Signal intensities of *ssr3189* were determined using Quantity One software (BIO-RAD) and normalized to the signal of the 5S rRNA.

The PNK assays were carried out as described (Brenes-Álvarez et al., 2025), with modifications regarding the growth of the cyanobacterial strains. 400 mL of cultures from reporter strains expressing the 3xFLAG fusion proteins under the control of the P*_petE_* promoter were grown in BG11 supplemented with the appropriate antibiotic until they reached OD ∼0.6. After 24 h of cultivation, half of each culture (200 mL) was treated (1.6778 mJ/cm2) while gently shaking to induce RNA-protein crosslinking keeping the sample on ice (the remaining half was also placed on ice). Cells were pelleted by centrifugation at 3,270 x g for 10 min. The pellets were resuspended in NT-P buffer (50 mM NaH_2_PO_4_, 300 mM NaCl, 0.05% Tween, pH 8.0) supplemented with protease Inhibitor (cOmplete EDTA-free, Roche). Cells were lysed as described in a Precellys homogenizer (Bertin Technologies). Glass beads were removed by centrifugation at 1,500 x g for 2 min. Next, membranes were pelleted by centrifugation at 21,000 x g for 30 min at 4 °C. 2 mL of clarified lysate per sample (crosslinked and non-crosslinked) was incubated with 80 μL of anti-FLAG M2 magnetic beads (Sigma) prewashed 5 times in NT-P buffer. From this point on, all the washing steps were performed with 2 mL of buffer. The samples were incubated at 4 °C while gently rotating for 1 h. Beads were washed twice with NT-P buffer, then once with benzonase buffer (50 mM Tris-HCl, 1 mM MgCl_2_, pH 8). The beads were resuspended in 100 μL benzonase buffer supplemented with protease inhibitor and 25 U of benzonase nuclease (Sigma). The samples were incubated at 37 °C for 10 min, while shaking at 900 rpm. After this, the beads were washed once with NT-P buffer and twice with FastAP buffer (10 mM Tris-HCl, 5 mM MgCl_2_, 100 mM KCl, pH 8). The beads were resuspended in 150 μL of FastAP buffer supplemented with protease inhibitor and 2 U of Fast Alkaline Phosphatase (Thermo). Samples were incubated at 37 °C for 30 min, shaking at 900 rpm. The beads were washed once with NT-P buffer and twice with PNK buffer (50 mM Tris-HCl, 10 mM MgCl_2_, 0.1 mM spermidine, pH 8). The magnetic beads were resuspended in 100 μL of PNK buffer supplemented with protease inhibitor, 10 μCi of [γ-32P]-ATP and 1 μL of PNK (Thermo). Samples were incubated at 37 °C for 30 min, while shaking at 900 rpm. The beads were washed once with NT-P buffer. For the elution of the protein-RNA complexes, the beads were resuspended in 35 μL of TBS buffer (50 mM Tris-HCl, 150 mM NaCl, pH 7.5) supplemented with 0.5 μg/μL FLAG peptide (Sigma) and incubated at 4 °C for 15 min, while gently shaking at 900 rpm. After separation of beads to the supernatant using a magnetic rack, protein loading dye was added to supernatant (5x concentrated loading dye: 25 mM Tris-HCl pH 6.8, 25% glycerol, 10% SDS, 50 mM DTT and 0.05% bromophenol blue). After 5 min of denaturation at 95 °C, the samples were separated on 12% SDS-PAA gels and electroblotted onto nitrocellulose membranes (Amersham). Radioactive ink was added to the corners of the membrane to facilitate the overlay of the size markers. Radioactive signals were detected using a Typhoon FLA 9500 (GE Healthcare). After detection of the radioactive signal, the membranes were blocked in 5% skim milk in TBS-T for 2 h and subsequently used for Western blotting using anti-FLAG antibody as already described.

The RNA samples of two biological replicates each were hybridized to 8×60K microarrays (Agilent ID 075764) following published sample preparation and hybridization details (Voß and Hess, 2014). In short, 2 µg of DNase-treated RNA was used for Cy3 labeling (ULS Fluorescent Labeling Kit for Agilent Arrays, Kreatech). Microarray hybridization was performed with 600 ng Cy3-labeled RNA for 17 h at 65 °C. Microarray raw data were obtained in an Agilent microarray Scanner C, Modell G2505C using Agilent Scan Control and Feature Extraction software version 10.7.3.1. and were processed using the limma R package (version 3.52.4) (Ritchie et al., 2015). The data was normalized using the R-implementation in limma (Ritchie et al., 2015) and resulting *p* values from t-tests were corrected using the Benjamini and Hochberg (Benjamini and Hochberg, 1995) method to control the False Discovery Rate of differential expression. A |log_2_FC| ≥ 0.6 threshold and a *p* value ≤ 0.05 were considered to indicate a significant change in gene expression. The full dataset is accessible in the GEO database with the accession number GSE342216.

### Co-IP, LC-MS/MS Analyses and Data Processing

Cultures grown with the CellDeg cultivator system (CellDEG GmbH) in 100 mL of fresh water organism (FWO) medium with a final concentration of 10 mM sodium bicarbonate (NaHCO_3_) at a starting OD_750_ of 0.8. The lower chamber of CellDeg contained a bicarbonate/carbonate buffer consisting of potassium bicarbonate (KHCO_3_) and potassium carbonate (K_2_CO_3_). Expression was induced with 2 μM of Cu^2+^, for 24 h and before harvesting the cultures by centrifugation (5,000 × *g*, 4 °C, 10 min) after 24 h of P*_petE_*-*ssr3189*×3F overexpression, resuspended in FLAG buffer (50 mM HEPES-NaOH pH 7; 5 mM MgCl_2_; 25 mM CaCl_2_, 150 mM NaCl; 10% glycerol; 0.1% Tween-20) supplemented with Protease Inhibitor (cOmplete, Roche) and lysed in a Precellys homogenizer (Bertin Technologies). To solubilize membrane proteins, extracts were incubated for 1 h in the presence of 2% n-dodecyl β-D-maltoside in the dark at 4 °C. To remove cell debris and glass beads, the samples were centrifuged at 13,000 × g for 30 min at 4 °C. The cleared total cell lysate was subjected to co-IP using ANTI-FLAG M2 affinity agarose beads (Sigma) for 2 hours at 4 °C under gentle rotation. The beads were washed 6 times with 2 mL of FLAG-buffer using DynaMag™-2 magnetic rack (Thermo Fisher). Elution of bound proteins was performed with 250 µL FLAG-peptide (Sigma) solution (400 ng/µL) in TBS buffer at 4 °C for 30 min and gentle rotation. The supernatant was taken as elution fraction. Three independent replicates were prepared. Eluate fractions were measured for protein concentrations with the Coomassie Plus Bradford Assay (Thermo Fisher Scientific), and 5 µg of protein was used for preparation of the samples for MS measurements. Samples were prepared by adding 1% SDS and 5 mM DTT, followed by an incubation at 60 °C for 30 min with shaking at 1,000 rpm. in a ThermoMixer® (Eppendorf). Reduced disulfides were alkylated by adding 0.1 vol of 2-chloroacetamide (200 mM) and incubation for 15 min at 37 °C and shaking. The alkylation reaction was quenched by adding 5 mM of DTT. The protein clean-up and digestion was performed following the SP3 protocol (Hughes et al., 2019). To each sample, 100 μg of a 1:1 mixture of Sera-Mag SpeedBeads suspension (prepared to a 50 μg/μL, GE Healthcare®, cat. No. 45152105050250 and 65152105050250) was added. To induce binding of the proteins to the beads, 4 vol of 100% ethanol were added and incubated in a ThermoMixer® at 24 °C for 5 min at 1,000 rpm. Tubes were placed in a magnetic rack and the beads were washed 3 times with 180 μL of 80% ethanol. Beads were resuspended in 75 μl of 100 mM ammonium bicarbonate, pH 8 containing 0.24 μg of sequencing grade, modified trypsin (Promega), sonicated for 30 s in a water bath to disaggregate the beads and incubated for 18 h at 37 °C in a ThermoMixer® at 1,000 rpm. After digestion, the samples were centrifuged at 20,000 × g for 1 min at RT. The tubes were placed in the magnetic rack and the supernatant, containing the digested proteins, was transferred to a new tube. Samples were acidified by adding formic acid to a final concentration of approximately 2% to obtain a pH < 3. Next, AttractSPE Tips RPS T1 200 μL (Affinisep) were equilibrated with 50 μL methanol centrifuged for 2 min at 1,000 × g, followed by the addition of 50 μL 0.5% acetic acid and another centrifugation. The acidified samples were loaded onto the equilibrated tips and centrifuged. Samples were washed with 30 μL and 60 μL of 0.1% formic acid and 80% acetonitrile. Samples were eluted with 65 μL of freshly prepared 5% ammonium hydroxide and 60% acetonitrile in water. Peptides were eluted by centrifugation for 3 min at 500 × g into glass vials. The elution was dried in SpeedVac set at 30 °C and stored at – 20 °C. For LC-MS analysis, an UltiMateTM 3000 RSLCnano system was online coupled to a Q Exactive mass spectrometer (both Thermo Fisher Scientific, Dreieich, Germany). The peptide mixture was washed and preconcentrated on a PepMap C18 trapping column (Thermo Fisher Scientific) with a flow rate of 30 μL/min and peptides were separated on a μPAC™ C18 pillar array column (200 cm bed length, Thermo Fisher Scientific) using a binary buffer system consisting of A (0.1% formic acid) and B (86% acetonitrile, 0.1% formic acid). Peptides from pull down samples were eluted applying a gradient with a total duration of 1h with a flow rate of 1 µl/min. For total lysate samples the gradient was of 2 h total duration at a flow rate of 0.5 μL/min. For electrospray ionization of peptides, a Nanospray Flex ion source (Thermo Fisher Scientific) with a liquid junction PST-HV-NFU (MS Wil, The Netherlands) and a fused silica emitter (20 μm inner diameter, 360 μm outer diameter, MicrOmics Technologies, Spanish Fork, UT) with a source voltage of +1,800 V and an ion transfer tube temperature of 275 °C were used. Cycles of data-independent acquisition (DIA) consisted of: one overview spectrum (RF lens of 50%, normalized AGC target of 3E6, maximum injection time of 60 ms, *m/z* range of 385 to 1,050, resolution of 70,000) recorded in profile mode followed by MS2 fragment spectra (RF lens of 50%, normalized AGC target of 1E6, maximum injection time of 60 ms) generated sequentially by higher-energy collision-induced dissociation (HCD) at a normalized energy of 26% in a precursor *m/z* range from 400 to 1000 in 50 windows of 12.5 *m/z* isolation width overlapping by 0.25 *m/z* and recorded at a resolution of 35,000 in profile mode.

MS raw data were converted to mzML format using the ProteoWizard software, version 3.0.21229 (Chambers et al., 2012). For protein and peptide identification and quantification a library free search was performed using DIA-NN version 1.8.1 (Demichev et al., 2020) against the organism-specific Uniprot database (release 2023_03, *Synechocystis* 6803, taxid 1148), complemented with the sequences of the bait protein with a part of the FLAG tag, restricted to the first three amino acids up to the first trypsin cleavage site, 3xFLAG and GFP. Trypsin was set as protease with 2 allowed missed cleavages. Carbamidomethylation of cysteine was selected as fixed modification, N-terminal excision of methionine and oxidation of methionine were selected as variable modifications with a maximum number of 2. For precursor ion generation, peptides ranging from 6 to 35 amino acids, precursor charge from 2 to 4 precursor *m/z* range from 350 to 1,100 and fragment ion *m/z* range from 200 to 2,000 were used. Further parameters were: match between runs, unrelated runs, search in double-pass mode, deactivated heuristic protein inference, robust LC quantification and no cross-run normalization. The protein group quantities output file from DIA-NN was used and all proteins with non-zero label-free quantification (LFQ) intensities in 3 out of 3 replicates in one group were selected for further analysis. Missing values were imputed in two steps: proteins with 3 missing values in one group (missing completely) were subjected to left-censored imputation using the MinProb method, followed by sequential imputation using the R package impSeqRob (Verboven et al., 2007) for proteins with ≥1 remaining missing values. For differential abundance analysis, log_10_ transformed LFQ intensities were subjected to a moderated t-test using the R package limma (Ritchie et al., 2015) with correction of p-values for multiple testing (Benjamini and Hochberg, 1995).

### SDS‒PAGE, Native Gel Electrophoresis and Western Blotting

Proteins were mixed with denaturing loading dye (5× concentrated: 250 mM Tris-HCl pH 6.8, 25% glycerol, 10% SDS, 500 mM DTT, and 0.05% bromophenol blue G-250). The protein samples were separated by 12% SDS‒PAGE using the PageRuler^TM^ molecular weight marker (Thermo Fisher). To run samples under nonreducing conditions, a native loading dye (5× concentrated: 30% glycerol, 0.05% bromophenol blue G-250, 150 mM Tris-HCl pH 7.5) was used, and boiling was omitted. To make proteins visible, the gel was stained with Coomassie dye. The gel was incubated for 30 min under slight shaking, covered with Instant Blue Coomassie Protein Stain (Abcam) and washed overnight with ddH_2_O. For immunoblot analysis, separated proteins were transferred to nitrocellulose blotting membrane (Amersham^TM^ Protan, pore size 0.45 µm). Membranes were blocked overnight at 4 °C with 5% milk powder in TBS-T, washed three times (10 min), and subsequently probed with monoclonal ANTI-FLAG^®^ M2-Peroxidase (HRP) antibody (Sigma–Aldrich, A8592 at a dilution of 1:5000) in TBS-T for 1 h at room temperature in the dark and washed again three times (10 min). All washing steps were performed with TBS-T (20 mM Tris pH 7.6, 150 mM NaCl, 0.1% (v/v) Tween-20) at room temperature. Signals were detected with WesternBright^TM^ ECL-spray (Advansta) on a chemiluminescence imager system (Fusion SL, Vilber Lourmat) and subsequently visualized using FUSION-CAP (Vilber Lourmat) and Quantity One software (BIO-RAD) as previously described (Baumgartner et al., 2016; Krauspe et al., 2021). For the Western blot against GlnA, the antiserum was used in a 1:15,000 dilution and incubated overnight at 4 °C. The secondary anti-rabbit antiserum (Sigma-Aldrich A8275) was used in a 1:10,000 dilution and incubated at room temperature for 1 h.

### Extraction and Metabolite Analysis

Wild type, Δ*ssr3189* and Δ*ssr3189::ssr3189*×3F complementation strain were cultivated in Erlenmeyer flasks in BG11 medium with Na_2_CO_3_ (0.38 mM final concentration) in triplicates without antibiotics. At the indicated time points, samples of 5 vol*OD were collected and snap frozen in liquid nitrogen. For extraction, 630 µL of methanol containing carnitine (internal standard, 1 mg per extraction) was added, mixed for 1 min, and incubated for 5 min in a sonic water bath. After another 15 min of shaking at room temperature, 400 µL of chloroform was added, and the sample was incubated at 37 °C for 5 min. Next, 800 µL of ROTISOLV LC‒MS grade H_2_O (Roth) was added, and the sample was mixed thoroughly. Precipitation was enabled by storage at-20 °C for at least 2 h, and phase separation was subsequently achieved by centrifugation for 5 min at room temperature (16,800 × *g*). The upper polar phase was collected and dried in a speed vac for 30 min followed by lyophilization overnight. Absolute metabolite contents were quantified on an HPLC‒MS-8050 system (Shimadzu) as described (Reinholdt et al., 2019). Dried samples were dissolved in 400 μL LC‒MS grade water and filtered through 0.2-mm filters (Omnifix-F, Braun, Germany), and 4 μL of the cleared supernatant was separated on a pentafluorophenylpropyl column (Supelco Discovery HS FS, 3 mm, 150 3 2.1 mm). The compounds were identified and quantified using the multiple reaction monitoring (MRM) values given in the LC‒MS/MS method package and the LabSolutions software package (Shimadzu). Authentic standard substances (Merck) at various concentrations were used for calibration, and peak areas were normalized to signals of the internal standard (carnitine). The data were further normalized to the OD_750_ measured for each sample. The raw data and estimations as absolute values in ng per mL per OD_750_ are given in **Supplemental Dataset 5**, with all the statistical evaluations.

### Determination of Enzymatic Activities Glutamine synthetase assay

Cultures were started with 10 mL of *Synechocystis* 6803 culture in BG11 medium, with an initial chlorophyll concentration of 0.4 µg/mL, in the absence of antibiotics. Cells were harvested by centrifugation after 7 d of culture, with chlorophyll concentrations of 10-15 µg/mL. Samples were centrifuged at 16,000 x g for 5 min at 4 °C. The supernatant was carefully removed and the precipitate corresponding to 10 mL of culture was resuspended in 4 mL of 50 mM Tris-HCl, pH 7.5. Cells were broken with a French press, twice at 1500 psi. One hundred µL from the samples obtained after the French press were used to determine GS transferase activity, as previously described (El Alaoui et al., 2001), during 30 min at 37 °C. The reaction mixture contained: 100 mM glutamine,10 mM sodium hydroxilamine, 50 µM manganese chloride,10 µM ADP and 50 mM sodium arseniate in 0.2 M MOPS, pH 7. GS activity was determined by triplicate for each of the samples, by using two dilutions of each extract (1/50 and 1/100). One unit of the activity is the amount of enzyme that transforms 1 mmol of substrate min^-1^. Protein concentration was determined using the Bio-Rad Protein Assay kit, based on the method described (Bradford, 1976). Chlorophyll concentration was determined as previously described (Mackinney, 1941). Results shown correspond to the average of three biological replicates; error bars correspond to the standard deviations.

### Enolase assay

Wild-type *Synechocystis* 6803 and Δ*ssr3189* mutant cells were disrupted by three passages through a French press (1,500 psi). Cell-free extracts were obtained by ultracentrifugation (60,000 × *g,* 4 °C) for 30 minutes and the resulting soluble fraction was used for enolase activity measurements. Enolase activity was determined using a continuous coupled spectrophotometric assay. Phosphoenolpyruvate formed from 2-phosphoglycerate was converted to pyruvate by pyruvate kinase (PK), followed by further reduction to lactate by lactate dehydrogenase (LDH), coupled to the oxidation of NADH to NAD⁺. Enzyme activity was monitored by measuring the decrease in absorbance at 340 nm resulting from NADH oxidation.

Standard assays (200 μL final volume) contained 50 mM Tris-HCl, pH 7.8, 10 mM 2-phosphoglycerate, 5 mM ADP, 10 mM MgCl₂, 0.7 mM NADH, 2 U PK (rabbit muscle; Merck, Darmstadt, Germany), 2 U LDH (rabbit muscle; Merck, Darmstadt, Germany) and 80 µg of soluble protein. Reactions were performed at 30 °C in 96-well microplates (BRANDplates^®^, BRAND, Wertheim, Germany), and the decrease in absorbance at 340 nm was recorded using an Infinite M200 plate reader (Tecan Group AG, Männedorf, Switzerland). NADH concentration was quantified against a standard curve (0 - 0.7 mM). Negative control reactions were performed in the absence of either the cell extract or 2-phosphoglycerate. One milliunit (mU) of enolase activity was defined as the formation of 1 nmol phosphoenolpyruvate, equivalent to the oxidation of 1 nmol NADH, per minute.

## DATA, MATERIALS, AND SOFTWARE AVAILABILITY

The transcriptomic (microarray) datasets are accessible in the GEO database with the accession number GSE342216. The mass spectrometry proteomics data have been deposited to the ProteomeXchange Consortium via the PRIDE (Perez-Riverol et al., 2022) partner repository with the dataset identifier PXD081927.

## ACKNOWLEDGMENTS

We are very grateful to Isabel Muro-Pastor and Francisco J, Florencio (both Sevilla, Spain) for providing GlnA antisera; Marcus Ziemann (Freiburg) for the support in bioinformatic analysis and helpful tools, and Annegret Wilde (Freiburg) for access to the spectrophotometer. The work of MYG was supported by the Intramural Research Program of the National Institutes of Health. The contributions of the NIH authors are considered Works of the United States Government. The findings and conclusions presented in this paper are those of the authors and do not necessarily reflect the views of the NIH or the U.S. Department of Health and Human Services.

## AUTHOR CONTRIBUTIONS

WRH, AK, MBR and MBA designed the project. AK, with input from NA, generated the mutant strains. AK, MBR, MBA and WRH performed the bioinformatic analysis, MG defined COG5974. AK and NA provided samples for microarrays which were performed by VR. NA performed RNA-related experiments and analyses. Metabolite extraction was performed by NA, ST and MH carried out the metabolic measurements and analyses. FD, BK and PH carried out MS-based proteomic analyses. Glutamine synthetase assays were performed by ALL and JMGF, enolase assays by RSO and BS. All other experiments were performed by AK and NA. AK, NA and WRH wrote the manuscript with input from all authors.

