## Supplement for "Deletion of the gene for a cyanobacterial ribosome-associated protein affects the carbon/nitrogen metabolism"

*Co-sharing first authors

### Supplemental Material

**Supplementary Datasets:** p. 2
**Supplementary Tables:** p. 4
**Supplementary Figures:** p. 8
**Supplementary References:** p. 25

**Supplementary Datasets**

**Supplemental Dataset 1. The 665 identified homologs of *ssr3189* in cyanobacteria.** The columns in the first sheet give the respective protein ID, followed by the NCBI taxonomy ID, the aa sequence, taxonomic details and comments. The second sheet provides details for the sequences used for the alignment shown in **Figure 1C**. This dataset extends **Figure 1** and **Figure S1.**

-- See separate Excel file –

**Supplemental Dataset 2. Members of COG 5794 (Clusters of Orthologous Genes).** This dataset extends **Figure 1** and **Figure S1.**

-- See separate Excel file –

**Supplemental Dataset 3. Measurements of growth curves and absorption spectra of the wild type and Δ*ssr3189* at standard conditions, and calculation of the growth rate and doubling time.** Sheet one: growth details; sheet two: absorption spectra raw data. This dataset extends **Figure 2A and 2B.**

-- See separate Excel file –

**Supplemental Dataset 4. Microarray data for the wild type and Δ*ssr3189* at standard conditions and nitrogen depletion for 3h and 24h.** This dataset is organized in 5 sheets comparing the data for the Δ*ssr3189* mutant (KO) to wild type (WT) under standard conditions, or after 3 and 24h of nitrogen starvation for the Δ*ssr3189* mutant (KO) and wild type (WT). Only genes with statistically significant differences ($|\log_2 FC| \geq 0.6$ and a $p$ value $\leq 0.05$) are included. Each sheet gives the following data in separate columns: the ID of the transcriptional unit (TU) as previously defined (Kopf et al., 2014), followed by the locus tag (Kaneko et al., 1996) and the gene IDs used in the microarray Agilent ID 075764. The following columns give, if available, the assigned gene names and a short description. The remaining three columns provide the numerical values for the calculated fold changes ($\log_2$FC), adjusted $p$ values and its -$\log_{10}$ values.

This dataset extends **Figure 4, 5, and Figure S5.**

-- See separate Excel file –

**Supplemental Dataset 5. LC-MS/MS data for 19 amino acids and several organic acids.** Retention time (r) was individually monitored with 2 standards (line 10 and 11) and the average peak area for 1 ng (line 13), which were used to determine the ng per sample. The carnitine standard (columns K, L, M, N) was used to generate the Factor IS (internal standard), which is implemented in the normalization of the peak area for each sample.

-- See separate Excel file –

**Supplemental Dataset 6. Proteins enriched together with Ssr3189 by co-IP, identified by mass spectrometry.** The proteins enriched in the Ssr3189 co-IP (PD_ssr3189) and a sfGFP mock co-IP (PD_Ctrl) were identified, as well as in the respective protein lysates from the cultures expressing Ssr31893xFLAG (CE_ssr3189) or sfGFP (CE_Ctrl), each in triplicates 1—3. This Table extends **Figure 8** and **Table 1.**

-- See separate Excel file –

### Supplementary Tables

**Supplemental Table S1. Glutamine synthetase activity and *t*-test.** This table extends **Figure 6B.** The assays were performed in triplicates numbered 1–3 with two biological replicates of wild type (WT1 and WT2), and triplicates of 3 independent clones of the Δ*ssr3189* mutant labeled c1, c2 and c3.

| **Cell extract** | **GS activity (U/mg protein)** |
|---|---|
| **WT1_1** | **0.898** |
| **WT1_2** | **0.886** |
| **WT1_3** | **0.920** |
| **WT2_1** | **0.787** |
| **WT2_2** | **0.868** |
| **WT2_3** | **0.871** |
| **Δ*ssr3189* c1_1** | **1.288** |
| **Δ*ssr3189* c1_2** | **1.207** |
| **Δ*ssr3189* c1_3** | **1.195** |
| **Δ*ssr3189* c2_1** | **1.044** |
| **Δ*ssr3189* c2_2** | **1.048** |
| **Δ*ssr3189* c2_3** | **0.998** |
| **Δ*ssr3189* c3_1** | **0.990** |
| **Δ*ssr3189* c3_2** | **1.01** |
| **Δ*ssr3189* c3_3** | **0.908** |

| | **Average** | **SD** | **P value** |
|---|---|---|---|
| **WT (n=6)** | **0.872** | **0.042** | **0.000935** |
| **Δ*ssr3189* (n=9)** | **1.076** | **0.117** | |

**Supplemental Table S2. Desoxyoligonucleotide primers used in this study**. Abbreviations: AQ, AQUA cloning (Beyer et al., 2015).

| **Name** | **Sequence (5‘ to 3‘)** | **Description** | **Purpose** |
|---|---|---|---|
| P_AK1 | GCAGGTCGACTTTCAGCCCACCGGAAATCTGCACC | AQ of Δ*ssr3189* construct to create flank1 (F1) with overlap to pUC19 plasmid backbone | Generation of the Δ*ssr3189* knock out strain with pUC19 plasmid |
| P_AK2 | CACTCTGTACTGTTTGTTCCTCATAAATATAGTTTACAG | AQ of Δ*ssr3189* construct to create flank1 (F1) with overlap to streptomycin resistance cassette (Strep$^{R}$) | |
| P_AK3 | AGGAACAAACAGTACAGAGTGATGTCAACG | AQ of Δ*ssr3189* to create the Strep$^{R}$ cassette with overlap to flank 1 | |
| P_AK4 | CTAAAAATAGATATTATCGTAGTTGCTCTCAG | AQ of Δ*ssr3189* to create the Strep$^{R}$ cassette with overlap to flank 2 | |
| P_AK5 | TACGATAATATCTATTTTTAGCGCTCCTAAGG | AQ of Δ*ssr3189* construct to create flank 2 (F2) with overlap to streptomycin resistance cassette (Strep$^{R}$) | |
| P_AK6 | GGATCCTCTAGTGGACGAGAACATCACCAAAATCC | AQ of Δ*ssr3189* construct to create flank 2 (F2) with overlap to streptomycin resistance cassette (Strep$^{R}$) | |
| P_AK7 | GGTGGGCTGAAAGTCGACCTGCAGGCATGCAAGC | AQ of Δ*ssr3189* to create the backbone of pUC19 with overlay to flank 1 (F1) | |
| P_AK8 | TTCTCGTCCACTAGAGGATCCCCGGGTACC | AQ of Δ*ssr3189* to create the backbone of pUC19 with overlay to flank 2 (F2) | |
| P_AK9 | TATTACGCCAGCTGGCGAAAGG | Sequencing the flank 1 of Δ*ssr3189* O construct in pUC19 | |
| P_AK10 | TGCCCTGCTTATTGAAGATGAGG | Sequencing the flank 1 of Δ*ssr3189* construct in pUC19 binding to streptomycin resistance cassette (Strep$^{R}$) | |
| P_AK11 | ATCCAACTACGACATTTCTCCAAGC | Sequencing the flank 2 of Δ*ssr3189* construct in pUC19 | |
| P_AK12 | CCAGGCTTTACACTTTATGCTTCCG | Sequencing the flank 2 of Δ*ssr3189* construct in pUC19 binding to streptomycin resistance cassette (Strep$^{R}$) | |
| P_AK13 | TCACAACAGTTTCCTCCGGG | Segregation check for *ssr3189* deletion | |
| P_AK14 | GGAGAAGTAGGAGCATTGTAGGC | Segregation check for *ssr3189* deletion | |
| P_AK15 | TTACAGATCCTCTAGAGtcgacccaacaaaccttagctaggg | Fragment for the native promotor of *ssr3189* with the overlap to the pVZ322 plasmid | Construction of Δ*ssr3189*::*ssr3189*×3F for complementation of Δ*ssr3189* in *Synechocystis* |
| P_AK16 | ccatcatgatctttataatcaacgtcgctggcggcaccg | Primer for the Ssr3189 ORF with 3xFlagTag | |
| P_AK17 | GATTATAAAGATCATGATGG | 3xFlagTag and 3'UTR for tagging small proteins | |
| P_AK18 | GATGTATGCTCTTCTGCTCCTGCAGTAATAAAAAACGCCCGGCG | 3xFlagTag and 3'UTR for tagging small proteins with overlap to the pVZ322 plasmid | |

| | | | |
|---|---|---|---|
| | GCAACCGAGCGAATAATTCCCA ACGAAGGCAAGC | | |
| P_AK22 | CTGACCTTGCCATCACGACT | Sequencing the *ssr3189* constructs on the pVZ322 plasmid | |
| P_AK23 | ATGAAAAATACCATGCTCAGAAA AGGC | Sequencing the *ssr3189* constructs on the pVZ322 plasmid | |
| P_AK17 | GATTATAAAGATCATGATGG | 3xFlagTag and 3'UTR for tagging small proteins | Construction of $P_{petE}$:: *ssr3189*×3F for ectopic expression in *Synechocystis* |
| P_AK18 | GATGTATGCTCTTCTGCTCCTG CAGTAATAAAAAACGCCCGGCG GCAACCGAGCGAATAATTCCCA ACGAAGGCAAGC | 3xFlagTag and 3'UTR for tagging small proteins with overlap to the pVZ322 plasmid | |
| P_AK19 | TTACAGATCCTCTAGAGTCGAC CTGGGCCTACTGGGCTATTC | Fragment for the *petE* promotor of Ssr3189 ORF with the overlap to the pVZ322 plasmid | |
| P_AK20 | GTTTTGCCATACTTCTTGGCGAT TGTATCTATAG | Fragment for the *petE* promotor of Ssr3189 ORF with the overlap to the translational start site | |
| P_AK21 | GCCAAGAAGTATGGCAAAACGT CGTAATTTAAAG | Primer for the Ssr3189 ORF at the translational start site with overlap to the *petE* promoter | |
| P_AK22 | CTGACCTTGCCATCACGACT | Sequencing the *ssr3189* constructs on the pVZ322 plasmid | |
| P_AK23 | ATGAAAAATACCATGCTCAGAAA AGGC | Sequencing the *ssr3189* constructs on the pVZ322 plasmid | |
| P_AK24 | TAATACGACTCACTATAGGTCTT TGGCCACCGGAATAG | Probe for *ssr3189* northern hybridization with T7 promoter | Probe for *ssr3189* northern hybridization |
| P_AK25 | gccaagaagtatggcaaaacgtcgtaatt taaag | Probe for *ssr3189* northern hybridization | |
| P_AK26 | TAATACGACTCACTATAGGGAG Gacttggcatcggactattgtgccgtggg | Probe for 5S rRNA northern hybridization with T7 promoter | Probe for 5S rRNA northern hybridization |
| P_AK27 | tcttggtgtctttagcgtcatgg | Probe for 5S rRNA northern hybridization | |

### Supplemental Table S3. Details of constructed plasmids.

| Plasmid name | Description | Reference |
|---|---|---|
| pUC19::Δ*ssr3189* | Plasmid used for homologous recombination of $Strep^R$ cassette into the *ssr3189* locus using as flanking sites the up- and downstream genomic regions flank 1 and flank 2 to create the deletion strain Δ*ssr3189*. | This study |
| pVZ322::$P_{ssr3189}$-*ssr3189*×3F | Self-replicating, conjugative plasmid for native *ssr3189* expression by $P_{ssr3189}$ promoter in *Synechocystis* 6803, $Km^R$ and $Gen^R$ to create the complementation strain Δ*ssr3189*::*ssr3189*×3F. | This study |
| pVZ322::$P_{petE}$-*ssr3189*×3F | Self-replicating, conjugative plasmid for copper-inducible *ssr3189* expression from $P_{petE}$ promoter in *Synechocystis* 6803, $Km^R$ and $Gen^R$ to create the overexpression strain $P_{petE}$-*ssr3189*×3F. | This study |

### Supplementary Figures

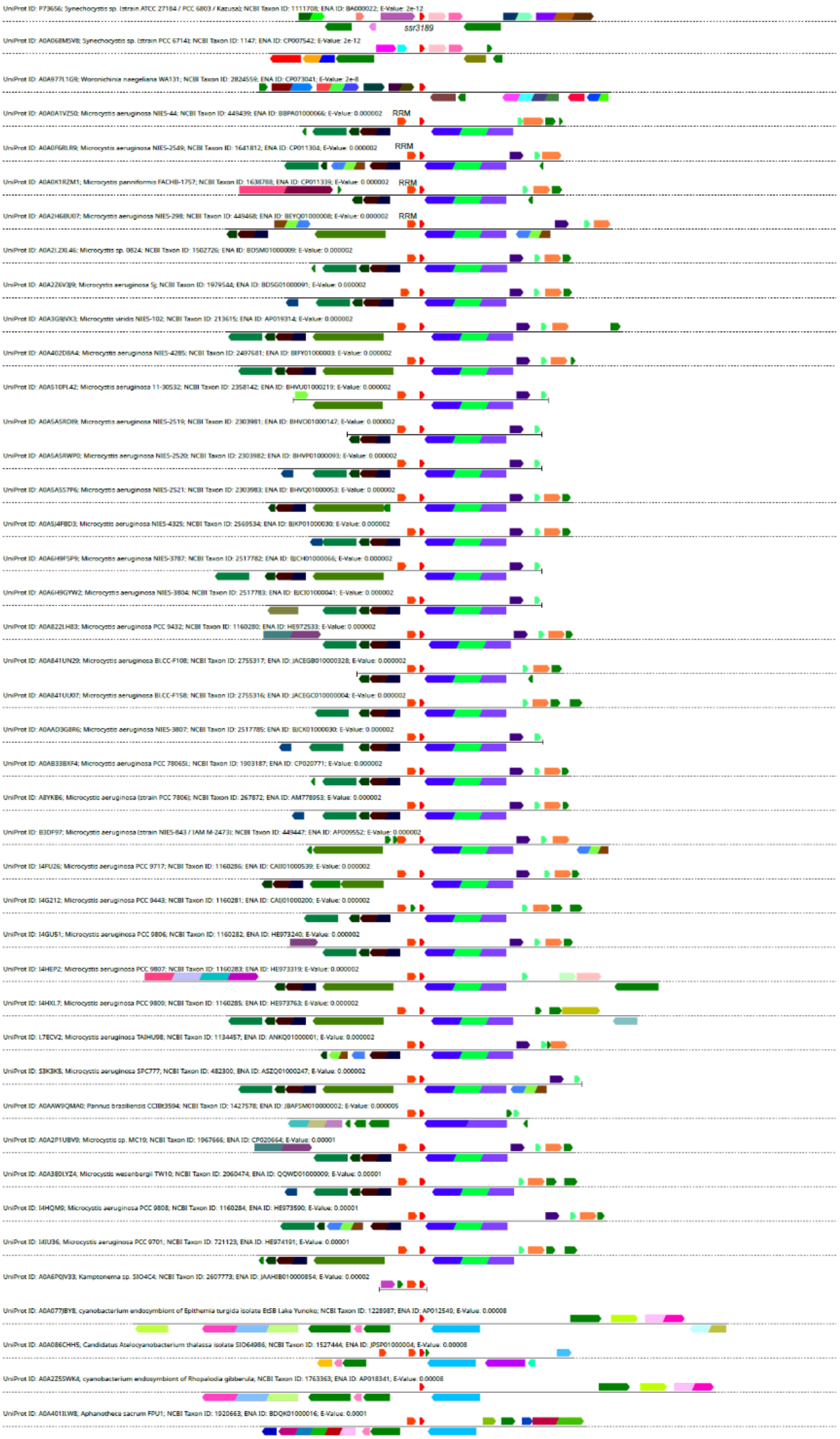

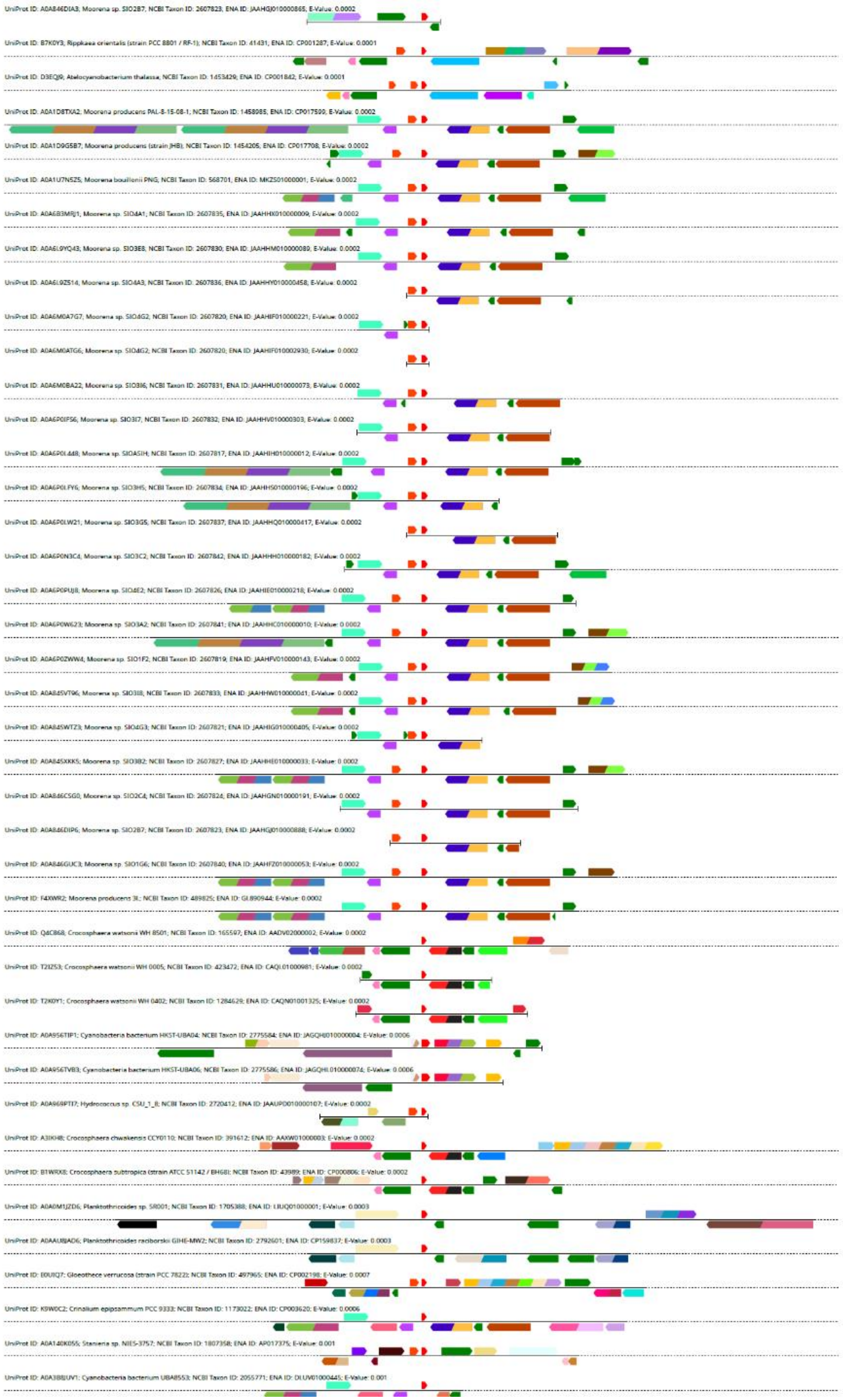
UniProt ID: A0A846DIA3; Moorena sp. SIO2B7; NCBI Taxon ID: 2607823; ENA ID: JAAHGJ010000865; E-Value: 0.0002
UniProt ID: B7K0Y3; Rippkaea orientalis (strain PCC 8801 / RF-1); NCBI Taxon ID: 41431; ENA ID: CP001287; E-Value: 0.0001
UniProt ID: D3EQJ9; Atelocyanobacterium thalassa; NCBI Taxon ID: 1453429; ENA ID: CP001842; E-Value: 0.0001
UniProt ID: A0A1D8TXA2; Moorena producens PAL-8-15-08-1; NCBI Taxon ID: 1458985; ENA ID: CP017599; E-Value: 0.0002
UniProt ID: A0A1D9G5B7; Moorena producens (strain JHB); NCBI Taxon ID: 1454205; ENA ID: CP017708; E-Value: 0.0002
UniProt ID: A0A1U7N5Z5; Moorena bouillonii PNG; NCBI Taxon ID: 568701; ENA ID: MKZS01000001; E-Value: 0.0002
UniProt ID: A0A6B3MRJ1; Moorena sp. SIO4A1; NCBI Taxon ID: 2607835; ENA ID: JAAHHX010000009; E-Value: 0.0002
UniProt ID: A0A6L9YQ43; Moorena sp. SIO3E8; NCBI Taxon ID: 2607830; ENA ID: JAAHHM010000089; E-Value: 0.0002
UniProt ID: A0A6L9Z514; Moorena sp. SIO4A3; NCBI Taxon ID: 2607836; ENA ID: JAAHHY010000458; E-Value: 0.0002
UniProt ID: A0A6M0A7G7; Moorena sp. SIO4G2; NCBI Taxon ID: 2607820; ENA ID: JAAHIF010000221; E-Value: 0.0002
UniProt ID: A0A6M0ATG6; Moorena sp. SIO4G2; NCBI Taxon ID: 2607820; ENA ID: JAAHIF010002930; E-Value: 0.0002
UniProt ID: A0A6M0BA22; Moorena sp. SIO3I6; NCBI Taxon ID: 2607831; ENA ID: JAAHHU010000073; E-Value: 0.0002
UniProt ID: A0A6P0IFS6; Moorena sp. SIO3I7; NCBI Taxon ID: 2607832; ENA ID: JAAHHV010000303; E-Value: 0.0002
UniProt ID: A0A6P0L448; Moorena sp. SIO5A1H; NCBI Taxon ID: 2607817; ENA ID: JAAHIH010000012; E-Value: 0.0002
UniProt ID: A0A6P0LFY6; Moorena sp. SIO3H5; NCBI Taxon ID: 2607834; ENA ID: JAAHHS010000196; E-Value: 0.0002
UniProt ID: A0A6P0LW21; Moorena sp. SIO3G5; NCBI Taxon ID: 2607837; ENA ID: JAAHHQ010000417; E-Value: 0.0002
UniProt ID: A0A6P0N3C4; Moorena sp. SIO3C2; NCBI Taxon ID: 2607842; ENA ID: JAAHHH010000182; E-Value: 0.0002
UniProt ID: A0A6P0PUJ8; Moorena sp. SIO4E2; NCBI Taxon ID: 2607826; ENA ID: JAAHIE010000218; E-Value: 0.0002
UniProt ID: A0A6P0W623; Moorena sp. SIO3A2; NCBI Taxon ID: 2607841; ENA ID: JAAHHC010000010; E-Value: 0.0002
UniProt ID: A0A6P0ZWW4; Moorena sp. SIO1F2; NCBI Taxon ID: 2607819; ENA ID: JAAHFV010000143; E-Value: 0.0002
UniProt ID: A0A845VT96; Moorena sp. SIO3I8; NCBI Taxon ID: 2607833; ENA ID: JAAHHW010000041; E-Value: 0.0002
UniProt ID: A0A845WTZ3; Moorena sp. SIO4G3; NCBI Taxon ID: 2607821; ENA ID: JAAHIG010000405; E-Value: 0.0002
UniProt ID: A0A845XKK5; Moorena sp. SIO3B2; NCBI Taxon ID: 2607827; ENA ID: JAAHHE010000033; E-Value: 0.0002
UniProt ID: A0A846CSG0; Moorena sp. SIO2C4; NCBI Taxon ID: 2607824; ENA ID: JAAHGN010000191; E-Value: 0.0002
UniProt ID: A0A846DIP6; Moorena sp. SIO2B7; NCBI Taxon ID: 2607823; ENA ID: JAAHGJ010000888; E-Value: 0.0002
UniProt ID: A0A846GUC3; Moorena sp. SIO1G6; NCBI Taxon ID: 2607840; ENA ID: JAAHFZ010000053; E-Value: 0.0002
UniProt ID: F4XWR2; Moorena producens 3L; NCBI Taxon ID: 489825; ENA ID: GL890944; E-Value: 0.0002
UniProt ID: Q4C868; Crocosphaera watsonii WH 8501; NCBI Taxon ID: 165597; ENA ID: AADV02000002; E-Value: 0.0002
UniProt ID: T2IZ53; Crocosphaera watsonii WH 0005; NCBI Taxon ID: 423472; ENA ID: CAQL01000981; E-Value: 0.0002
UniProt ID: T2K0Y1; Crocosphaera watsonii WH 0402; NCBI Taxon ID: 1284629; ENA ID: CAQN01001325; E-Value: 0.0002
UniProt ID: A0A956TIP1; Cyanobacteria bacterium HKST-UBA04; NCBI Taxon ID: 2775584; ENA ID: JAGQHJ010000004; E-Value: 0.0006
UniProt ID: A0A956TVB3; Cyanobacteria bacterium HKST-UBA06; NCBI Taxon ID: 2775586; ENA ID: JAGQHL010000074; E-Value: 0.0006
UniProt ID: A0A969PTI7; Hydrococcus sp. C5U_1_8; NCBI Taxon ID: 2720412; ENA ID: JAAUPD010000107; E-Value: 0.0002
UniProt ID: A3IKH8; Crocosphaera chwakensis CCY0110; NCBI Taxon ID: 391612; ENA ID: AAXW01000003; E-Value: 0.0002
UniProt ID: B1WRX8; Crocosphaera subtropica (strain ATCC 51142 / BH68); NCBI Taxon ID: 43989; ENA ID: CP000806; E-Value: 0.0002
UniProt ID: A0A0M1JZD6; Planktothricoides sp. SR001; NCBI Taxon ID: 1705388; ENA ID: LIUQ01000001; E-Value: 0.0003
UniProt ID: A0AAU8JAD6; Planktothricoides raciborskii GIHE-MW2; NCBI Taxon ID: 2792601; ENA ID: CP159837; E-Value: 0.0003
UniProt ID: E0UIQ7; Gloeothece verrucosa (strain PCC 7822); NCBI Taxon ID: 497965; ENA ID: CP002198; E-Value: 0.0007
UniProt ID: K9W0C2; Crinalium epipsammum PCC 9333; NCBI Taxon ID: 1173022; ENA ID: CP003620; E-Value: 0.0006
UniProt ID: A0A140K055; Stanieria sp. NIES-3757; NCBI Taxon ID: 1807358; ENA ID: AP017375; E-Value: 0.001
UniProt ID: A0A3B8JUV1; Cyanobacteria bacterium UBA8553; NCBI Taxon ID: 2055771; ENA ID: DLUV01000445; E-Value: 0.001

**Figure S1. Synteny analysis of *ssr3189* and its homologs in cyanobacteria.** The analysis was performed using the EFI Genome Neighborhood Tool (Oberg et al., 2023).This figure is an extension to **Figure 1.**

Likely homologs are in identical colors (red, *ssr3189* and homologs; light red, homologs of the gene encoding RRM domain protein Rbp1; rose, *apaH*; purple, *trpS*).

**CYANOBACTERIA**

```
                         --hhhhHHHHHHHHHHHHHHHHhhhh-----------------------------
BAA17701.1          1    MAKRRNLKKEKAERNRAYARQFRSASRTQNRRYSGGQRQQGDSNGSSNNGAASDV   55 (55)
BAB75587.1          1    MAKRRNPKKEKALRNQAYARKFRKRTTTGRMQRRFQQRTPKSEEEEGAAAVDAE-   54 (54)
ABW25224.1          1    MAKRRNLKKEKALRNQAYARKFRKRSGSGRFGRRQYNSNRQESNSDNQNGDNQDS   55 (66)
BAC91068.1          1    MAKRRNQKKEKALRNRANARKYRKKQVTGRYGRRSAGPGGFEDNDTGAQAP----   51 (51)
CAE20761.1          1    MAKRRNLKKEKQERNRSYARKFKKRKLRNDGRGEGAGNGVTGTANNGGAAD-----  51 (51)
BAC09286.1          1    MAKRRNPKKEKALRNQAYARKFRKRSSGRYSRRLNTENRPETKTTETVEEMTDA-   54 (54)
```

**Cyanobacterial endosymbionts of eukaryotes**

```
ADB95749.1          1    MAKRRNLKKEKAERNQAYARQFRQKASRSFSRGRSYSNNRSDQEDMNAADTTNDN   55 (56)
BDA39275.1          1    MAKRRNLKKEKAERNQAYARQFRQKTSRSFSRGRSYSNNRSDQEDTNDDDTSTNN   55 (55)
ACB42705.1          1    MAKRRNLKKEKQERNRAYARKFKKRKLRNDGRGEGAGNGVTGTANNGGAAD----   51 (51)
BBL86025.1          1    MAKRRNLKKEKQERNRAYARKFKKRKLRNDGRGEGAGNGVTGTANNGGAAD----   51 (51)
```

**EUKARYOTES**

**Green algae**

```
XRA98681.1         37    MACKRNLKKEKRTRNRINSMKYKKKPPPRFARRNNDVTEAKNDDWESPFLTDENG   91(131)
XRB13187.1         38    MACKRNLKKEKRTRNRINSMKYKKKPPPRFARRNNDVTEAKNDDWESPFLTDENG   92(132)
CAG9465386.1       36    MAKRRNAKKEKRQRNKEAARKYKKASNKGRRNFGGPETENQGAGWVSPFLYTPEE   90 (96)
```

**Red algae**

```
BAM79225.1         51    MRQRRDLREEKRLRNLEYARLHRKRVPRGPPGRRVNLSGNAAASAAERESVYSLY  105(140)
BHE93320.1         39    MRCRRDLKAEKRVRNSDFARKHRKRPVRRTGFVRPTPAAAAAAATARAEQAEKDE   93(168)
EME27828.1         50    MRSRRDLKKEKKLRNLEFAKLHRKRKERKRKVDQETLRVMDDNTFIKEVFSAEEN  104(107)
OSX70122.1         62    MRCRRNLKLEKRARNRANGRTFKKSKPRFFNRRSAVDTTTDKDTEFLSSIFNVLT  116(141)
CAO5933118.1       45    MRCRRDLKKEKSLRNLEFARSHRKKVIRRFNRRAAQEATQNEDNEYLSSIYGTMR   99(115)
KAA8499032.1       37    MACRRNLKREKICRNMAFARSHRPKSSEQRGNWRVAKKANEDSDDAYLNEIFSTL   91(110)
```

**Brown algae**

```
CAM9117422.1       55    MSCRVNAKKEKRKRNRDNMRKFTKRGTSKRKIMRGERQERAAEAEANFMAQIFHT  109(122)
CAM9182861.1       46    MSCRVNAKKEKRTRNRLNMRKFNKRGTSRKKILRVERQERAAEDEAEFLAKVFHS  100(113)
CAM9562096.1       53    MASRPNAKKEKRQRNRDNMRKFTKKGTSRKKVMRVQRQERAAEAESDFMAQVFHT  107(121)
CAM9461189.1       54    MGCRVNAKKEKRQRNRDNMRKFNKRGVSRKKVMRVERQERAAEAESNFMAQVFHT  108(121)
CAM9860172.1       26    MSCRVNAKKEKRQRNRDNMRKFNKKGTSRRRILKVARQERAAEVESNFMAEVFHS   80 (94)
CAM9568188.1       54    MGCRVNAKKEKRQRNRDNMRKFNKRGVSRKKVMRVERQQKAAEAEAEYVAQIFHS  108(121)
```

**Cryptomonads**

```
EKX39983.1         74    MSCRRNLKKEKRLRNEICARQYRKPKSKFRGSRPGRNARAMEEDADSEWLSQIYG  128(141)
KAJ1488946.1       76    MGCRRNLKKEKRLRNETCARKFRKPKPRFGGRPPMGAKKEDNGDSLWLEQIYGQH  130(163)
```

**Diatoms**

```
EEC50430.1         37    FACRSNAKKEKIKRNRENMRKFKTTGRKGTTRRKLLKKAQASKARQEESEFISKC   91(106)
EJK59850.1         43    DACRINAKQEKRKRNRENMRKFKKGGRKGTSKKKLMRKAASSAQRQVENEFIAKC   96(110)
OEU09409.1         91    FGCRVNAKKEKIKRNRENMRKFSTPGKRGMSRRKILKKAQASEARQKENEFITKC  145(158)
GAB7155838.1       37    FACRTNAKKEKRLRNRDNMRKFQKRGKTSRKKLMRKVASSEQRQLENEFVAKCFT   91(102)
```

**Haptophytes**

```
EOD19200.1         44    MACRSNLKKEKRLRNRVNAFRYKKGGFGKKRFSNNRFEDRAVAAAKATEDAEFLS   98(120)
GAC3484911.1       54    MACRHNSKKEKRLRNRINAFRFKKQTYGFNKFNNFADRNAQKANEQADGEWLAQV  108(128)
KAG8465933.1       49    MACRTNTKREKFFRNLENAKRFRKASWKPAFKGRGGPRSNDSTPESKEDAALMAT  103(127)
KAL1512177.1       48    MACRTNLKKEKRLRNRVNAFRFKKGGPVRFVRPGQYTPEDAKKASEDAEFYSLVY  102(123)
```

**Pelagophytes**

```
KAH8065851.1       39    EACRRNTKKEKRQRNQENMRKFKRGAAPAQRGAGGKKKSLSRKKLTLKAQSAKEK   93(120)
CAH0366694.1       41    EACRRNTKKEKRQRNQENMRKFKKGAPPTARGGGGKKKTMSRKKLTLKAQAAVEK   95(122)
```

**Yellow-green algae**

```
KAG5177122.1       43    MACRNNAKKQKAKRNQDNMRKFRKRGTSKKKIVRGKRQQTAREKEADFMAKLYQL   97(104)
```

**Figure S2. Sequence conservation of Ssr3189 homologs in distinct phylogenetic lineages.**

The top line shows the secondary structure prediction from AlphaFold, H indicates prediction of an α-helix with pLDDT > 70, h indicates less confident prediction (50 < pLDDT < 70), the dashes indicate a predicted unstructured region. The numbers indicate the positions of the start and the end of the alignment, the total protein lengths are shown in parentheses. Conserved residues are shown in bold: negatively (D, E) and positively (K, R) charged residues are shown in red and blue, respectively; conserved asparagine residues are in purple; conserved hydrophobic residues are shaded yellow. Note that the homologs from nuclear-encoded genes in eukaryotic algae are N-terminally longer, containing targeting sequences. This figure extends **Figure 1C.**

The proteins are listed under their GenBank accession numbers and linked to the respective entries in the NCBI protein database. The source organisms and gene names (top to bottom) are as follows:

**Cyanobacteria**: *Synechocystis* sp. PCC 6803, Ssr3189; *Nostoc* sp. PCC 7120, Asl3888; *Acaryochloris marina*, AM1_0138; *Gloeobacter violaceus*, gsl3127; *Prochlorococcus marinus* MIT 9313, PMT_0586; *Thermosynechococcus vestitus*, tsr1734.

**Cyanobacterial endosymbionts of eukaryotes:** *Cand*. Atelocyanobacterium thalassa (UCYN-A), UCYN_10740; *Braarudosphaera bigelowii*, nitroplast, CPARK_000011400; *Paulinella chromatophora*, chromatophore, PCC_0256; *Paulinella micropora*, chromatophore PMYN1_Chma213.

**Green algae:** *Pycnococcus provasolii*, NFJ02_05g123800; *Pseudoscourfieldia marina*, RI054_05g30240; *Pedinophyceae* sp. YPF701, YPF701_LOCUS5433.

**Red algae:** *Cyanidioschyzon merolae*, CYME_CMD165C; *Cyanidium caldarium* CdcalHa5_g4312.t1; *Galdieria sulphuraria*, Gasu_46500; *Porphyra umbilicalis*, BU14_0892s0001; *Delisea pulchra*, DPULC_LOCUS6866; *Porphyridium purpureum,* FVE85_6617.

**Brown algae:** *Desmarestia herbacea* PHAE_DHERF_3.8246.1; *Ectocarpus siliculosus* PHAE_ESILM_5.12546.1; *Fucus distichus*, PHAE_FDIS_17611.5239.1; *Halopteris paniculata* PHAE_HPAN_2394.6477.1; *Saccorhiza polyschides*, PHAE_SPOLM_11270.1157.1; *Sphacelaria rigidula* PHAE_SRIGF_7802.1.1.

**Cryptomonads:** *Guillardia theta*, GUITHDRAFT_154277; *Baffinella frigidus*, T484DRAFT_1938160.

**Diatoms:** *Phaeodactylum tricornutum*, PHATRDRAFT_43428; *Thalassiosira oceanica*, THAOC_19885; *Fragilariopsis cylindrus*, FRACYDRAFT_264135; *Chaetoceros neogracilis*, CNG_g02669.

**Haptophytes:** *Emiliania huxleyi*, EMIHUDRAFT_458783; *Chrysotila haptonemofera*, CHAPT_001836000; *Diacronema lutheri*, KFE25_005503, *Prymnesium parvum*, AB1Y20_005443.

**Pelagophytes:** *Aureococcus anophagefferens*, JL722_235; *Pelagomonas calceolata*, PECAL_1P32000.

**Yellow-green algae:** *Tribonema minus*, JKP88DRAFT_227007.

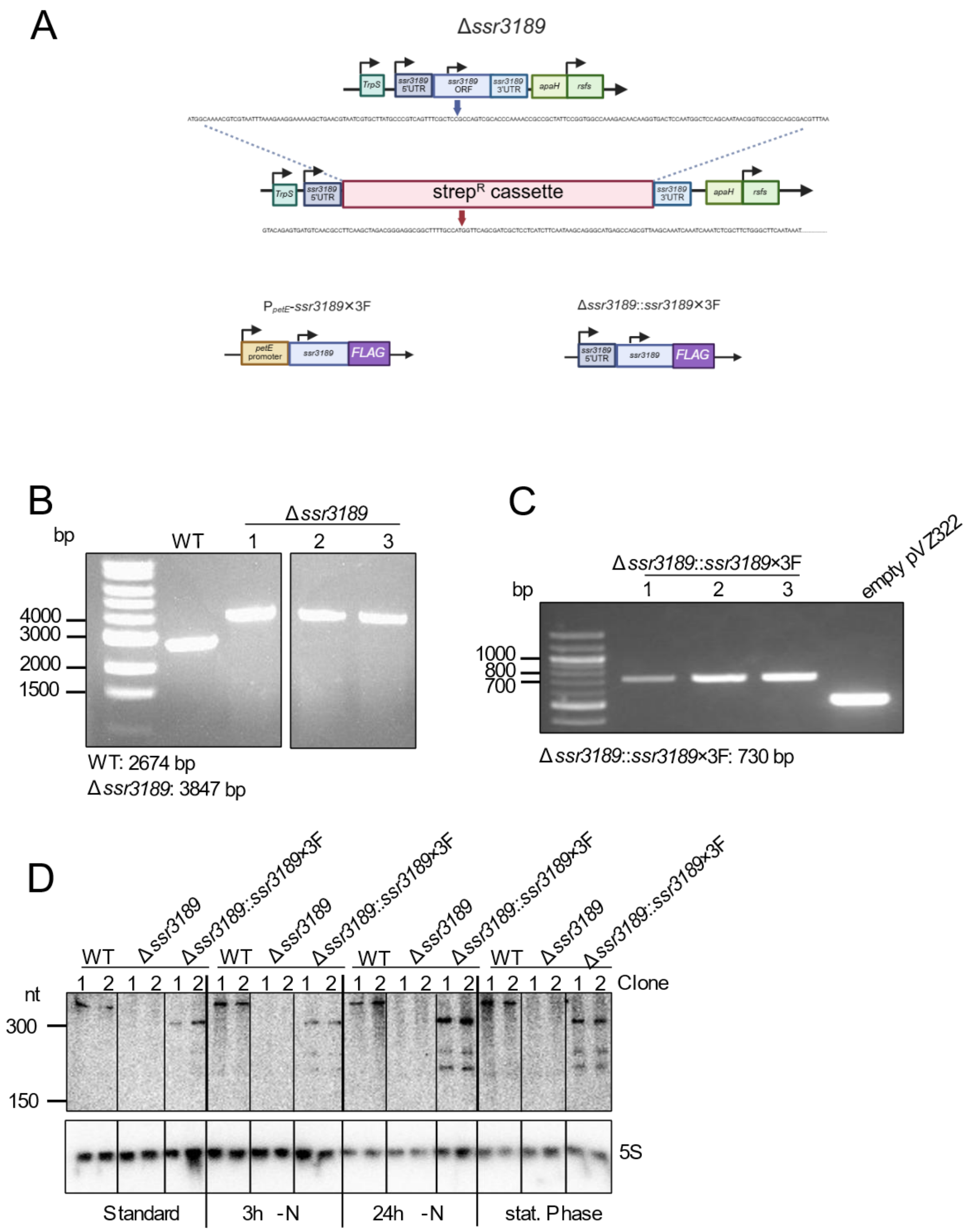


**Figure S3. Generation of Δ*ssr3189* and *ssr3189*-expressing strains. A.** The *ssr3189* locus in the wild type and in the Δ*ssr3189* deletion strain, as well as of a pVZ322 plasmid derivative harboring a *ssr3189* gene copy under the control of the $Cu^{2+}$-inducible $P_{petE}$ promoter in the *ssr3189* overexpression strain ($P_{petE}$-*ssr3189*×3F), and the complementation strain (Δ*ssr3189*::*ssr3189*×3F) under the control of its native promoter. The *ssr3189* gene was previously assigned to the transcriptional unit (TU)1276 (Kopf et al., 2014) extending from position 1244330 to 2217039 on the forward strand of the *Synechocystis* 6803 chromosome (GenBank accession no. NC_000911). In the Δ*ssr3189* strain, the gene was replaced by a streptomycin resistance cassette (*aadA*) (Hollingshead and Vapnek, 1985) ($Strep^R$) via homologous recombination. The plasmid enabling ectopic *ssr3189* expression was introduced into *Synechocystis* 6803 wild type and Δ*ssr3189* strain. **B.** PCR verification of the genotypes of independently obtained Δ*ssr3189* strains. Three clones were tested each, using primers P_AK13/P_AK14 (primer sequences in **Table S1**). **C.** PCR verification of the genotypes of independently obtained Δ*ssr3189*::*ssr3189*×3F complementation strains. Three clones were tested each, using the primers P_AK22/ P_AK23 (primer sequences in **Table S1**; bp, base pairs). **D.** Time course expression and inducibility of *ssr3189* mRNA in nitrogen deprivation after 3 and 24 h, in standard (OD ~0.8) and stationary phase were tested (OD ~4.0). The *ssr3189* transcript was detected by a $^{32}P$-labeled single-stranded RNA probe. The membrane was rehybridized to a 5S rRNA probe as a loading control. The RiboRuler Low Range RNA ladder (Thermo Fisher Scientific) was used as the molecular mass standard. This figure extends **Figure 2**.

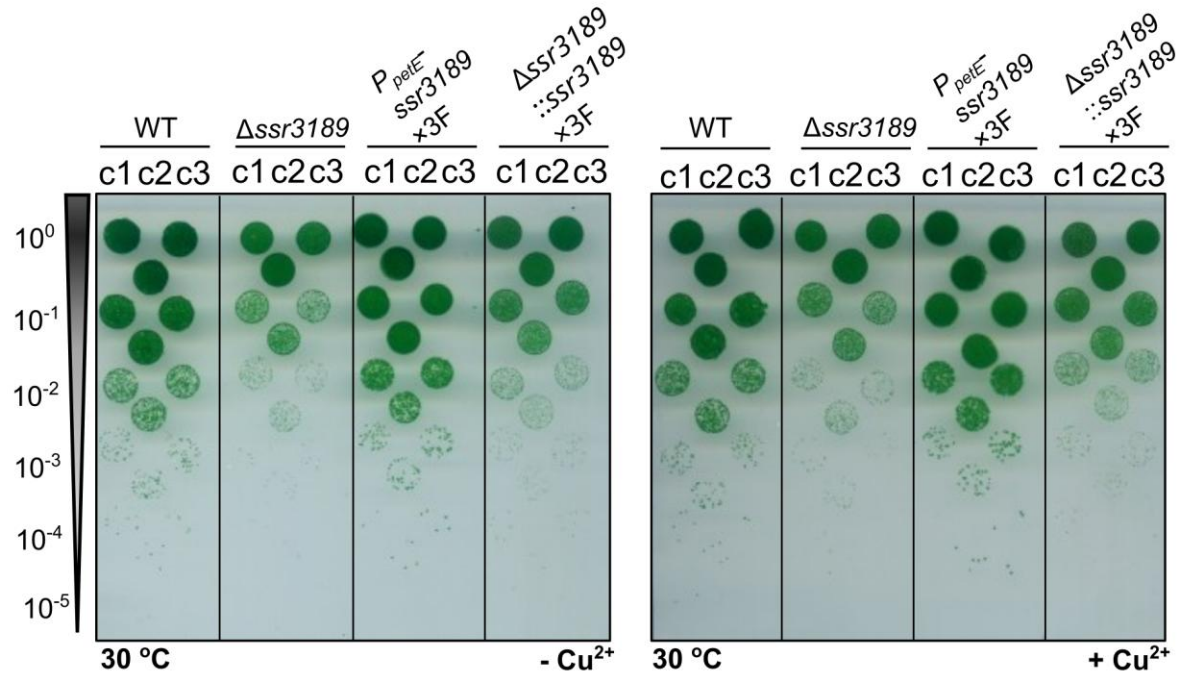


**Figure S4. Phenotypical differences between *ssr3189* mutant strains on plates.** Left panel: Drop dilution assay on solid medium comparing biological triplicates, c1 to c3, of the *Synechocystis* 6803 wild type (WT), the *ssr3189* deletion mutant Δ*ssr3189,* the complementation and overexpression strains Δ*ssr3189*::*ssr3189*×3F and $P_{petE}$-*ssr3189*×3F, at standard conditions, but without copper. Strains were pre-cultivated in liquid BG11 medium, $P_{petE}$-*ssr3189*×3F was pre-cultivated in BG11 without $CuSO_4$, under constant light at 30 °C. The indicated different dilutions were spotted on agar plates. Right panel: BG11 medium containing 0.3 µM $CuSO_4$. Plates were photographed after 5 days of incubation. This figure extends **Figure 3.**

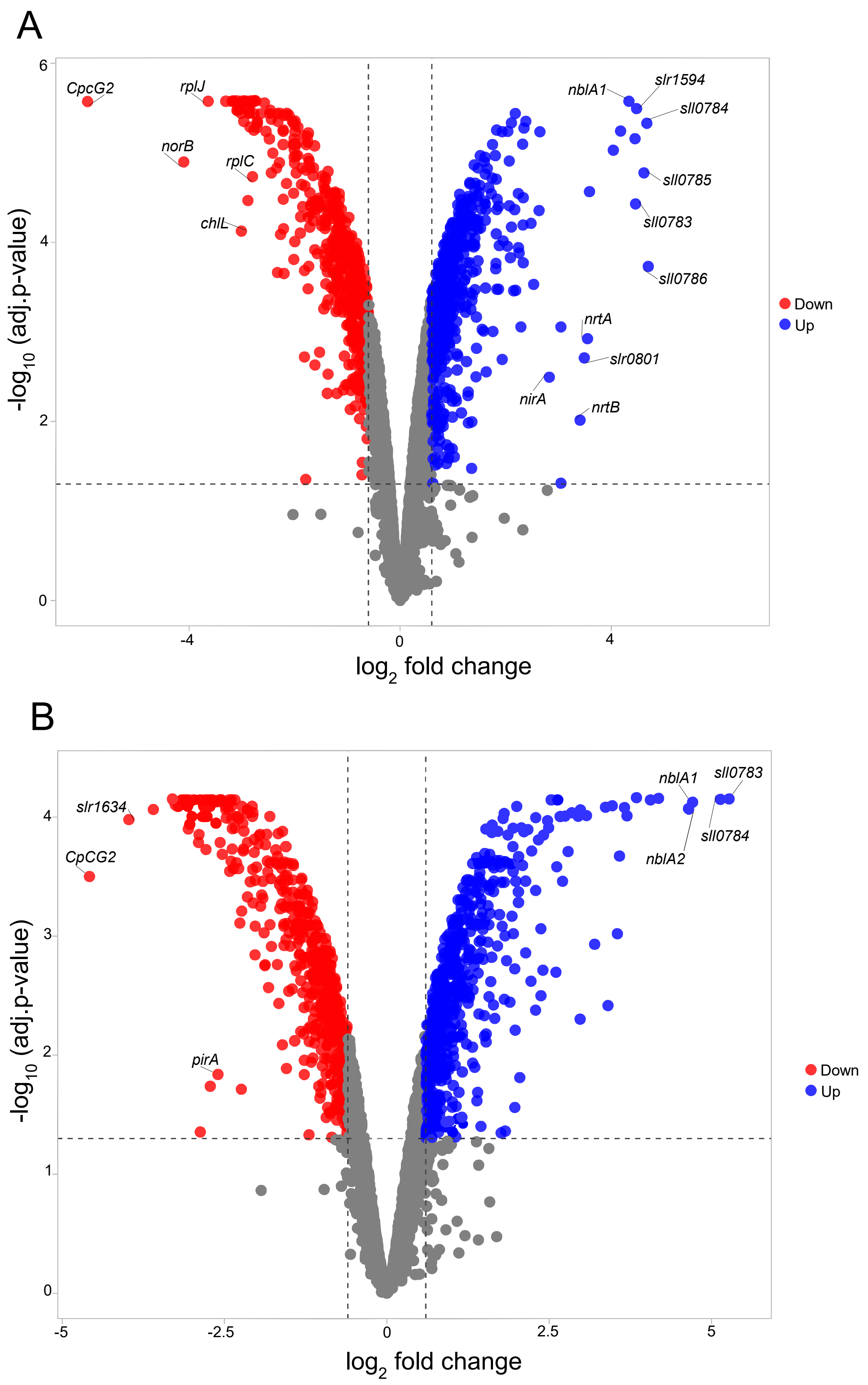
A
CpcG2
rplJ
norB
rplC
chlL
nblA1
slr1594
sll0784
sll0785
sll0783
sll0786
nrtA
slr0801
nirA
nrtB
Down
Up
-log10 (adj.p-value)
log2 fold change
B
slr1634
CpCG2
pirA
nblA1
sll0783
sll0784
nblA2
Down
Up
-log10 (adj.p-value)
log2 fold change

**Figure S5. Transcriptomic response to nitrogen starvation in *Synechocystis* 6803. A.** Volcano plot of microarray analysis of wild type in nitrogen-depleted media for 24 hours compared to wild type in nitrogen-depleted media for 3 hours. **B.** Volcano plot of microarray analysis of Δ*ssr3189* in nitrogen-depleted media for 24 hours compared to Δ*ssr3189* in nitrogen-depleted media for 3 hours. Upregulated genes are shown in blue and downregulated genes in red. Thresholds were set at $|log_2FC| \geq 0.6$ and a p value of $\leq 0.05$. This figure extends **Figure 5.**

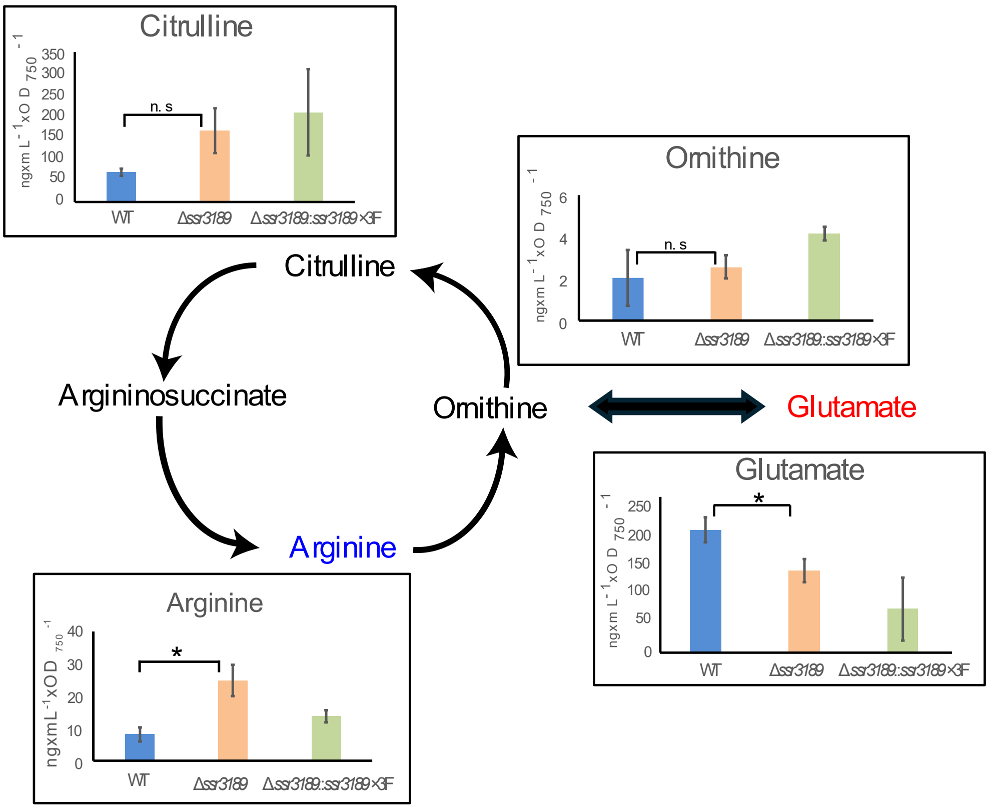


**Figure S6. Metabolites in the cyanobacterial ornithine-ammonia cycle.** Comparison of metabolites content (ng × $mL^{-1}$ × $OD_{750}^{-1}$) in wild type, Δ*ssr3189,* and Δ*ssr3189*::*ssr3189*×3F at standard conditions. Blue text indicates metabolites with significantly higher abundance, and red letters indicate metabolites with significantly lower abundance in Δ*ssr3189* relative to WT. Significance was calculated with two-sample *t* test with unequal variance (Welch's *t* test, n.s., not significant, *; $P < 0.05$).

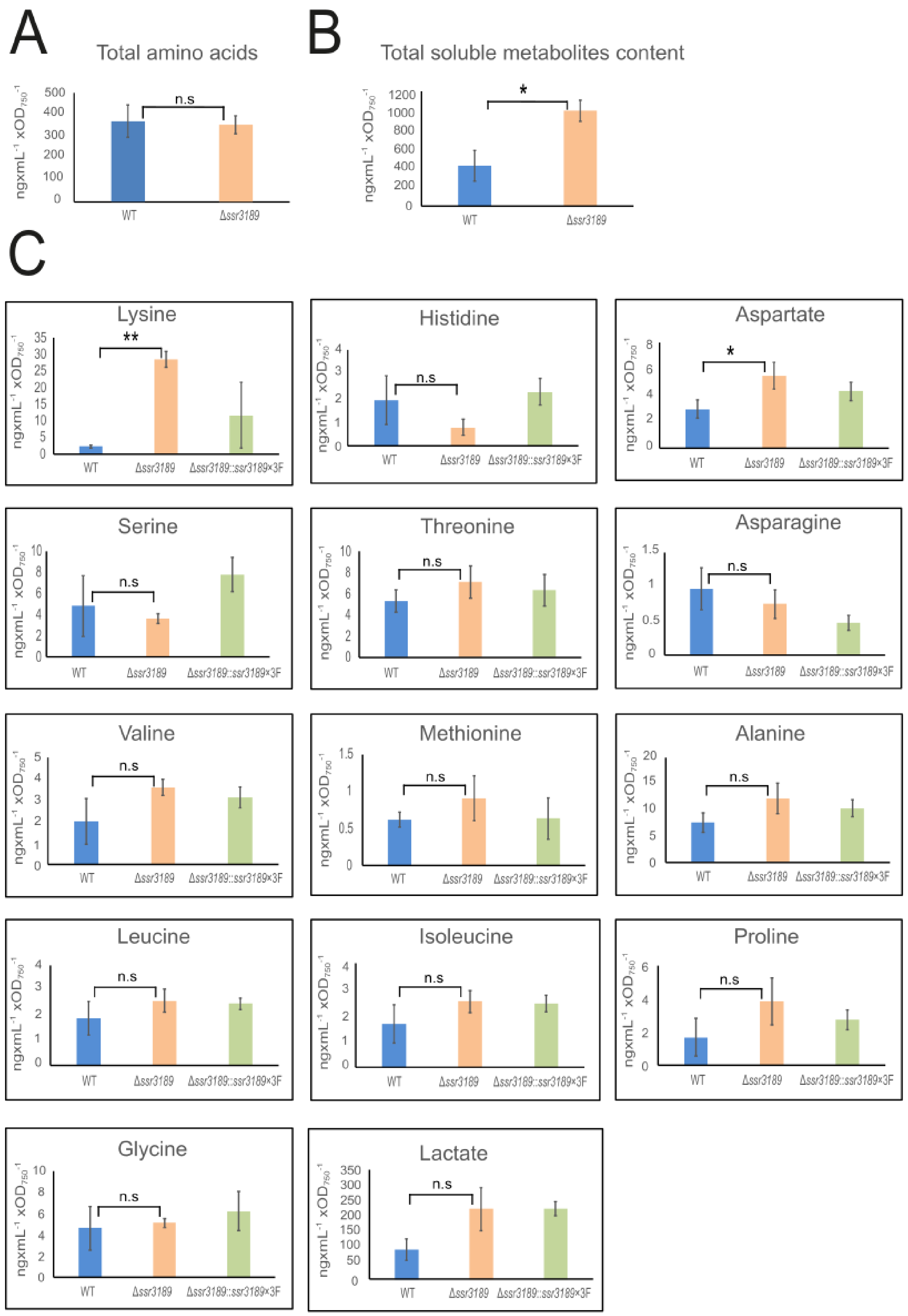


**Figure S7. Measurement of amino acids and metabolites. A.** Total soluble amino acid content and **B.** Total soluble metabolite content (amino acids and organic acids)

of the wild type and Δ*ssr3189*. **C.** Measured amino acids and lactate in the wild type, Δ*ssr3189* and Δ*ssr3189*:: Δ*ssr3189*×3F. Significance was calculated with two-sample *t* test with unequal variance (Welch's *t* test, n.s; not significant, *; $P < 0.05$, **; $P < 0.01$).

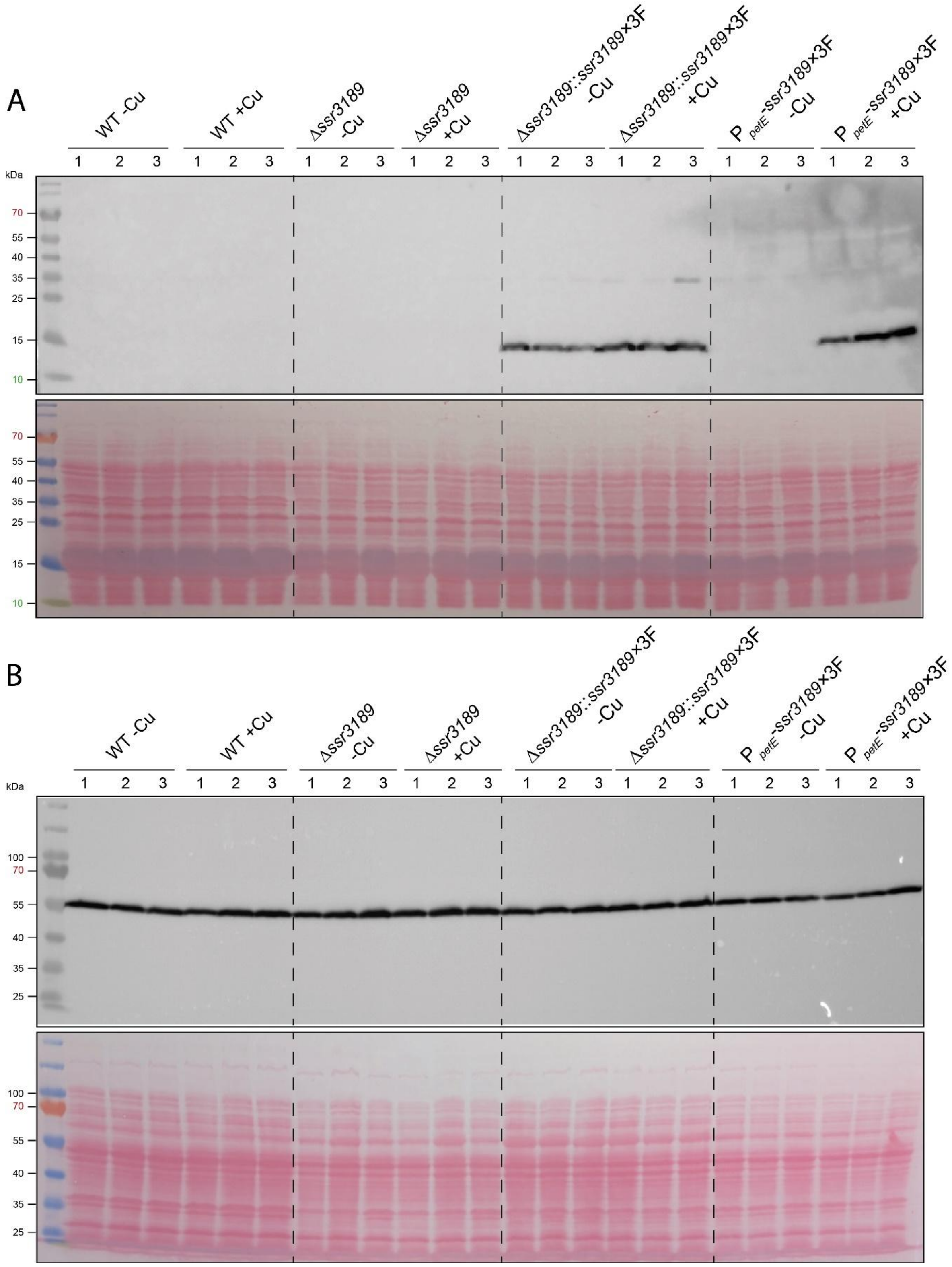


**Figure S8**. **Western blot validation of Ssr3189 expression and glutamine synthetase (GlnA) abundance. A.** Western Blot verification of the 3xFLAG-tagged Ssr3189 protein expressed from plasmid pVZ322 under control of the native promoter ($P_{ssr3189}$) or $P_{petE}$ promoter induced by copper addition to a final concentration of 1.25 µM (+Cu) for 24 h. Samples without induction served as negative controls (-Cu). The labels ‘1’ - ‘3’ refer to biological replicates. In the gel, 5 µg protein was loaded per lane, separated on a 15% SDS-PAA gel and incubated with M2 anti-FLAG HRP antibody. The lower panel shows the ponceau red-stained membrane. **B.** Shows the same samples as in A but separated on a 10% SDS-PAA gel and incubated with anti-GlnA antibody. This figure extends **Figure 6**.

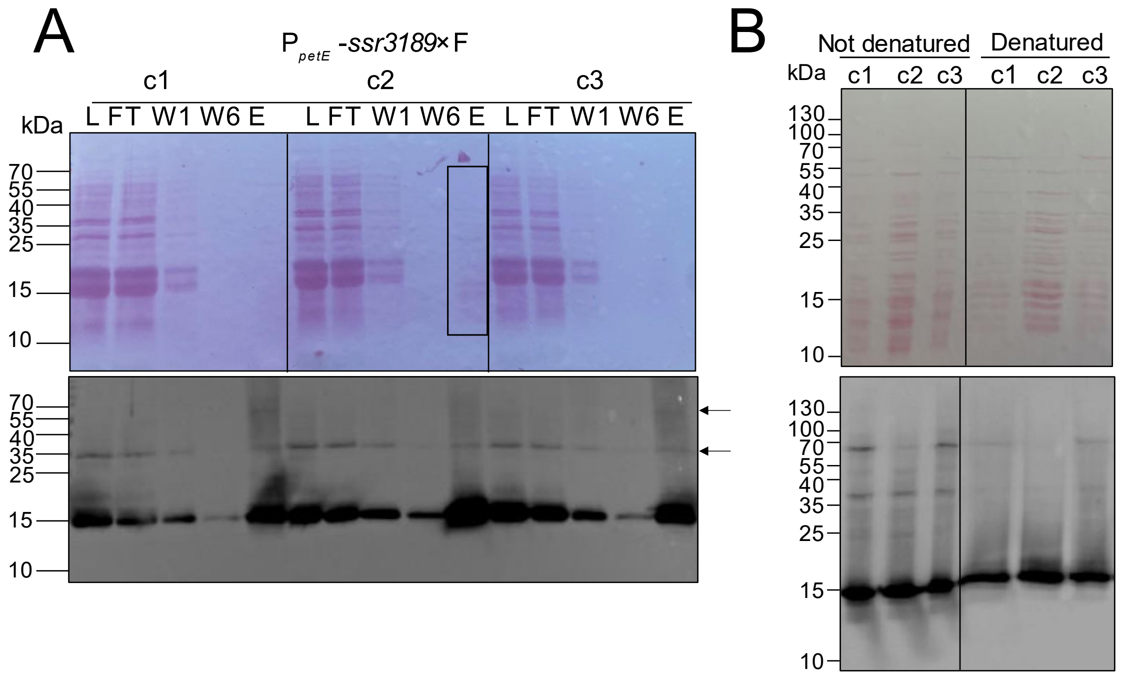


**Figure S9. Co-immunoprecipitation pull-down experiment revealed numerous interaction partners or multimerization potential of P$_{petE}$-*ssr3189*×3F. A.** Ssr3189 pull-down analysis of three biological replicates. Upper panel: Ponceau-stained membrane. Fractions are numbered and labeled CE, cell extract; FT, flow-through; W, wash; E, elution. Lower panel: Immunological identification of the lower band as Ssr3189. **B.** Western blot analysis of the three elution samples from panel (A) under denaturing or non-denaturing conditions. The samples were denatured by boiling in SDS loading buffer (Denatured), or not treated and loaded with non-denaturing loading buffer (Not denatured). This figure extends **Figure 8**.

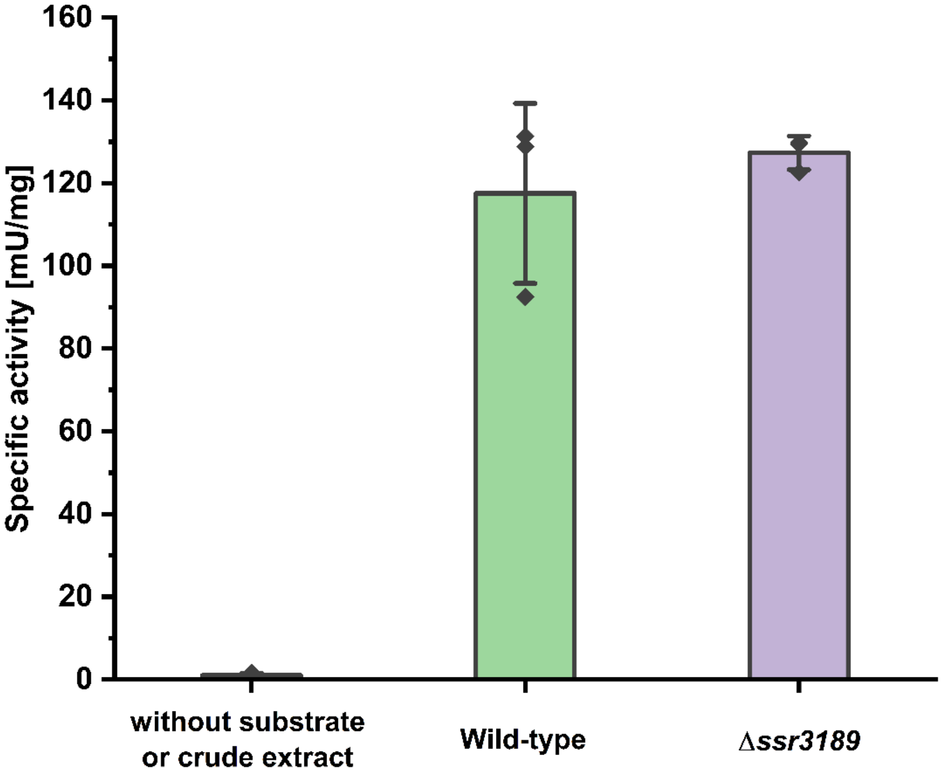


**Figure S10. Measurement of enolase activity in crude extracts of the wild type and the Δ*ssr3189* mutant.** Data are presented as mean ± SD from three technical replicates (n = 3).
